# Measurement and prediction of unmixing-dependent spreading in spectral flow cytometry panels

**DOI:** 10.1101/2025.04.17.649396

**Authors:** Peter L. Mage, Andrew J. Konecny, Florian Mair

## Abstract

Advances in spectral cytometry instrumentation and fluorescent reagents have led to the possibility of ultra-high-parameter panels exceeding 50 colors. However, panel size is limited in practice by unmixing-dependent spreading (UDS), a mathematical phenomenon which leads to a progressive deterioration of unmixed signal-to-noise ratios in panels that contain fluorochrome combinations with significant spectral overlap. Choosing spectrally compatible sets of fluorochromes that avoid UDS is a complex and labor-intensive task involving substantial trial-and-error experimentation. Here, we provide a detailed explanation of UDS and practical strategies for handling UDS in large spectral panels. We describe the empirical hallmarks of UDS, demonstrate how to quantify its impact, and dissect its underlying mathematical cause in terms of spectral collinearity. We present novel computational metrics that can be used to select optimal combinations of fluorochromes in a platform-agnostic fashion based on publicly available reference data, providing a general tool for spectral panel design.

## INTRODUCTION

The field of flow cytometry’s shift to the spectral paradigm^1–3^, along with advances in instrumentation and reagent design, have led to a rapid increase in the maximum plexity possible in a single flow cytometry experiment, with the simultaneous measurement of 50 fluorescent analytes now reported^4^. These high-parameter panels offer several advantages over combining separate lower complexity panels. First, for samples limited in size (*e*.*g*. pediatric blood samples) they maximize the number of cellular populations that can be analyzed. This is particularly relevant in the field of immunology, where it has become obvious that the interplay of several cellular populations, often at a specific tissue site, regulates an immune response in health and disease^5,6^. Second, combining all markers in a single staining allows identification of unexpected expression patterns that would otherwise be missed^7^.

However, expanding to higher-plexity panels is complicated by the fact that larger panels must inevitably use combinations of fluorochromes with high spectral overlap. Due to the physics of fluorescence detection and the mathematics of spectral unmixing, the signal-to-noise ratio of unmixed or compensated data is inextricably linked to the degree of spectral overlap among the fluorochromes being used. A well-known example of this tradeoff between panel size and measurement resolution is the phenomenon of “spillover spreading error” (SSE)^8^ that exists both in spectral and conventional cytometry. SSE occurs when photonic shot noise from emission of one fluorochrome leads to an increase in the variance of another spectrally overlapping fluorochrome’s compensated or unmixed signal. The impact of SSE on biological resolution must be managed through careful panel design^9–11^ and appropriate staining controls such as fluorescence-minus-one (FMO) samples^12,13^.

An additional data quality tradeoff has recently been observed in large spectral panels containing combinations of fluorochromes that are difficult to resolve from each other, wherein the variance of unmixed data increases irrespective of actual expression levels for a given marker. We have termed this effect “unmixing-dependent spreading” (UDS)^4^. While panel size was historically limited by available reagents and the number of detectors on a given instrument, panel size on spectral cytometers today is limited in practice by a progressive deterioration of the signal-to-noise ratio of unmixed data caused by UDS as the overall degree of spectral overlap in a panel increases. Therefore, predicting, diagnosing, and mitigating UDS in the context of a full panel is a key challenge when designing ever-larger panels without sacrificing biological resolution. To the best of our knowledge, no tools or platform-agnostic systematic workflows have been described to do so until now.

Existing spectral panel design tools have limited utility in diagnosing UDS. Empirical metrics based on measured data, exemplified by the Spillover Spreading Matrix (SSM)^8^, can reveal when SSE in unmixed data is worsened in the context of one panel compared to another, but fail to detect other manifestations of UDS. Moreover, these metrics require access to measured FCS data for all fluorochromes of interest, which may not be available early in the panel design process. Calculating these metrics is computationally intensive and requires bioinformatics expertise to re-unmix raw spectral data with multiple matrices for statistical analysis (a difficult workflow with most existing graphical-user-interface data analysis software). By contrast, predictive panel design metrics like cosine similarity (CS, sometimes called “similarity index”) and spectral matrix condition number (CN, sometimes called “panel complexity” or “complexity index”)^14,15^ can instead be calculated from spectral signatures alone, many of which are available in public reference databases such as commercial online spectrum viewers. While useful in some cases, these metrics cannot specify which fluorochromes cause UDS in a given panel. There remains a need for metrics that quantitatively predict problematic combinations of fluorochromes in the context of a specific panel.

In this manuscript we provide new tools for understanding, diagnosing, and avoiding UDS. First, we present a formal definition of UDS along with three hallmarks describing its typical representation in spectral cytometry data. Second, we describe an empirical approach for quantifying UDS in spectral data with specific metrics that reveal the degree of UDS for a given combination of fluorochromes. Third, we derive a practical mathematical tool called the “Hotspot Matrix” that predicts UDS based on spectral signatures alone and can thus be used to diagnose, predict, and avoid UDS as part of the panel design process, even prior to running any experiment. Fourth and finally, we provide a rigorous mathematical explanation of the underlying mechanism of UDS based on the concept of collinearity in multiple regression.

The approach described here was motivated by the authors’ development of 50-color flow cytometry in OMIP-102^4^. We hope that it will be of value to the wider flow cytometry community as a time- and cost-saving *in silico* tool that can be applied early in the panel design process, even before any reagents have been ordered or samples have been run.

## MATERIALS AND METHODS

### Sample preparation and data acquisition

For single-stain analysis, human PBMCs and whole blood were procured and stained with anti-CD4 antibodies conjugated to various fluorochromes. Data were acquired on a 5-laser BD FACSDiscover™ S8, 5-laser BD FACSymphony™ A5 SE, 5-laser Cytek® Aurora, or 7-laser Sony ID7000™ at default manufacturer-recommended detector settings unless otherwise noted. Detailed sample preparation, panel staining, and data acquisition protocols are provided in the Supplementary Materials.

### Data analysis and unmixing

Single-stain FCS data were processed in Python using custom scripts. Briefly, FCS files corresponding to each fluorochrome of interest were imported as *pandas* DataFrames using the *fcsparser* library. Lymphocyte scatter gates were defined using an automatic density-based gating algorithm. Next, CD4+ and CD4-lymphocyte populations were identified automatically based on fluorescence intensity using k-means clustering through the *scikit-learn* library^16^. Automated scatter and intensity segmentation results were confirmed via manual inspection. Each fluorochrome’s single-stain spectral signature was calculated by taking the difference of CD4+ and CD4-median fluorescence intensity (MFI) in each detection channel, setting any negative values to 0, and dividing by the MFI of the maximum-intensity detector so that the spectral signature spans the interval [0,1]. A single autofluorescence (AF) spectrum was defined based on unstained lymphocytes.

Spectral unmixing via ordinary-least squares (OLS) was performed in Python unless otherwise noted. AF was always included as a single color in the spectral matrices used for unmixing unless otherwise noted. Statistical and numerical linear algebra calculations were performed using the *numpy* library. Full-panel data was gated in FlowJo™ v10 (BD) and plotted in Python. Unmixed data was plotted using arcsinh (asinh) axis scaling with varying cofactors as appropriate.

### Statistical analysis

Population-level statistics including MFI and robust standard deviation (rSD) were calculated for each unmixed parameter for the CD4+ and CD4-populations in each fluorochrome recording. For unstained samples, statistics were calculated on all events in the lymphocyte scatter gate. rSD for an unmixed parameter *f* was calculated using the equation

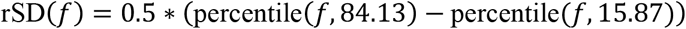

per the documentation for FlowJo™ v10 (BD).

Unmixing spreading error (USE) was quantified by taking the ratio (“rSD ratio”) of an unstained population’s rSD for a given parameter *f* when unmixed with a full-panel matrix *X* compared to unmixing with a single-color matrix *X*_1_:

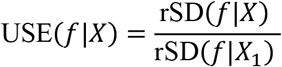

SSE and the SSM were calculated for unmixed fluorochrome parameters according to their original specification^8^.

### Unmixing metric calculations

Cosine similarity (CS or “similarity”) is defined as the cosine of the angle between the vectors defined by two fluorochromes’ spectral signatures^14,17^ (Supplementary Figure 1). CS was calculated for each pair of fluorochromes by first normalizing each fluorochrome’s spectral signature to have unit length (by dividing by its Euclidean norm) and subsequently taking the dot product of the two resulting normalized spectral signatures. The “similarity matrix” *S* containing all pairwise CS corresponding to the fluorochromes in a given spectral matrix *X* can also be calculated by normalizing each column of *X* to have unit length as above, and subsequently using the resulting normalized matrix *X*_*N*_ to calculate the matrix product 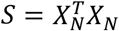. This is equivalent to an uncentered correlation matrix of the columns of *X*.

Condition number (CN or “complexity”) for a given spectral matrix was calculated by computing the singular value decomposition (SVD) of the spectral matrix and subsequently calculating the ratio of the largest singular value to the smallest singular value^18^. Following convention in the field, CN was always calculated using spectral signatures normalized to their maximum value.

A given panel’s Hotspot Matrix *H* (derived below) was calculated by finding the inverse matrix of that panel’s similarity matrix and subsequently taking the square root of each entry’s absolute value: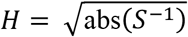. Spreading inflation factors (SIFs) are defined as the diagonal values of the Hotspot Matrix. The “SIF ratio” for a given fluorochrome was calculated as the ratio of that fluorochrome’s full-panel SIF to that fluorochrome’s single-color SIF (derived from the diagonal of the Hotspot Matrix for a spectral matrix containing only that color and AF, if AF was included in the full matrix).

## RESULTS

### Defining unmixing-dependent spreading (UDS) and its empirical hallmarks

The practical implications of UDS initially revealed themselves during testing of an early iteration of OMIP-102. We observed a substantial loss of resolution in unmixed data when expanding from a 20-color (20C) backbone panel to a 51C panel (Figure 1A) on a Sony ID7000™ cytometer. We initially ascribed this effect to SSE from added fluorochromes in the larger panel. However, we observed that the same raw data for the 20C backbone panel showed much higher spread across all measured cells when unmixed using the 51C spectral matrix compared to the 20C matrix (Figure 1A, center), suggesting that the spectral matrix itself had a profound impact on unmixed spread.

**Figure 1:**
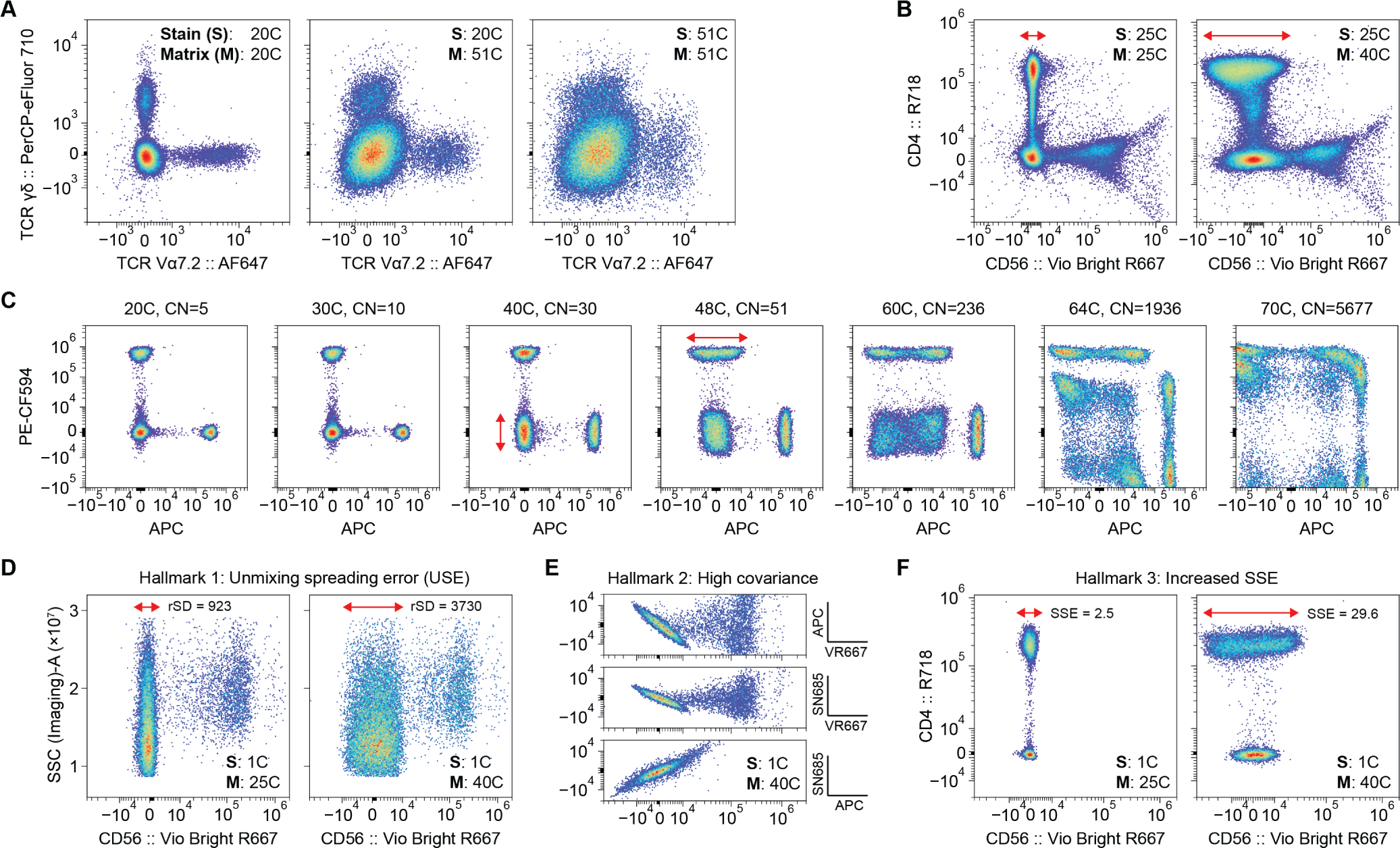
Observation and hallmarks of unmixing-dependent spreading (UDS). **(A)** Increased spreading error (SE) was observed when moving from a 20C backbone panel (left) to an expanded 51C spectral panel (right) when testing an early version of OMIP-102 on the Sony ID7000™. When the raw 20C backbone data was unmixed using the 51C matrix (middle), an increase in SE was still observed for the same raw data. Plots are pre-gated on CD14-CD3+ T cells. **(B)** Similar observations for an unrelated 25C backbone panel evaluated on the BD FACSDiscover™ S8, unmixed using a 25C matrix (left) or an expanded 40C matrix for a candidate panel design (right). Significant spillover spreading error (SSE) from R718 into Vio Bright R667 is observed only with the 40C spectral matrix (right). Cells shown are CD45+ lymphocytes. **(C)** UDS tends to increase for the same combinations of fluorochromes as panel size and condition number (CN) increase, as shown here with the same raw CD4 single-stain data unmixed in the context of panels of increasing size. These panels were designed with an algorithm which combines fluorochromes with minimal cosine similarity. **(D)** UDS Hallmark 1: Unmixing spreading error^4^ (USE) is an increase in SE independent of fluorochrome signal, as shown on the negative population of CD56:: Vio Bright R667 single-stain data unmixed with different matrices. **(E)** UDS Hallmark 2: “Tilted double-negative populations” with high positive or negative covariance, unrelated to biological expression patterns, are observed between certain pairs of fluorochromes in CD56:: Vio Bright R667 single-stain data unmixed with a 40C matrix (“VR667”: Vio Bright R667; “SN685”: Spark NIR 685). **(F)** UDS Hallmark 3: UDS can cause an increase in the severity of SSE between pairs of fluorochromes, even those that would not typically affect each other, shown for CD4:: R718 single-stain data. All matrices include autofluorescence (AF) as an unmixed parameter, with matrix size denoting the number of fluorochromes in the matrix exclusive of AF.

This phenomenon, which we termed UDS, was characterized by a noticeable spreading of the negative population in a given bivariate plot of unmixed parameters, often with a distinct “tilt” between the unmixed parameters. We noted that not all fluorochromes in the panel were affected to the same degree; many other unmixed parameters in the sample were unaffected (data not shown). We also observed UDS to impact SSE. When evaluating data from a 25C backbone panel unmixed with an expanded panel’s 40C matrix on the BD FACSDiscover™ S8 (Figure 1B), we found that UDS led to a dramatic increase in otherwise undetectable SSE from R718 into Vio Bright R667, illustrating the potentially unpredictable impact of UDS on panel performance.

To more clearly illustrate the impact of UDS, we examined a single pair of fluorochrome single-stained CD4 samples unmixed with matrices corresponding to panels of increasing size and condition number (Figure 1C). We observed that for the same underlying raw data, more complex unmixing matrices indeed led to an increase of variance in all measured cells. At 40C, background spreading in the APC parameter substantially increased; at 48C, SSE was observed to increase from PE-CF594 into APC; at 64C, the increase in spread due to UDS in both parameters nearly reached the level of the otherwise brightly resolved CD4 signal; and at 70C (on a 78-detector spectral cytometer) the data was essentially unusable.

Based on these observations, we propose the following definition of UDS: “a change in variance of an unmixed parameter when the same raw data is unmixed using a different spectral matrix containing a different set or number of fluorochrome spectral signatures.” As we will demonstrate below, UDS arises from the mathematics of the unmixing process itself and is determined by the spectral matrix. Unlike other sources of measurement variance in flow cytometry, it does not introduce additional random noise arising from some physical phenomenon. Rather, it propagates and amplifies measurement uncertainty that is already present in the raw data. It should be emphasized that UDS can occur regardless of whether a given fluorochrome is physically present in a sample. It depends solely on the presence or absence of problematic fluorochrome signatures in the matrix used for unmixing.

In our experience, UDS has three empirical hallmarks when observed in the context of spectral flow cytometry panels:

- **Hallmark 1:** Increased variance of unmixed parameters regardless of fluorochrome intensity, so-called “negative spread” or “unmixing spreading error” (USE)^4^ for its striking effect on the spread of unstained populations (Figure 1D). This unmixing spreading error affects all events in a sample, increasing the baseline degree of variance of all populations, but does not occur to the same degree across all fluorochromes. Because this hallmark affects all cells regardless of marker co-expression, USE can have a drastic impact on panel performance and the resolution obtainable for a given fluorochrome.
- **Hallmark 2:** High covariance between unmixed parameters, leading to the visual presentation of “tilted” or “diagonal” double-negative populations as an artifact unrelated to biological expression patterns (Figure 1E), similarly to what has been described for compensation-based cytometry data^19^.
- **Hallmark 3:** Increased magnitude of SSE between pairs of fluorochromes when unmixed with a full-panel matrix, including between pairs of fluorochromes that would not typically have high SSE (Figure 1F).

UDS may occur, in principle, in any multicolor flow cytometry experiment that utilizes compensation or spectral unmixing, and has in fact been observed in compensation-based experiments with closely overlapping fluorochromes^19^. In practice, however, we have observed it to occur most commonly in large spectral panels (40+ colors) in which several fluorochrome combinations with high similarity must inevitably be used. UDS may occur in smaller panels that contain highly overlapping fluorochromes; while such combinations are generally not preferable from a panel design perspective, they may still be chosen due to limited reagent availability, cost considerations, assay constraints, or expediency.

Importantly, UDS may occur regardless of the specific cytometer in use; we have observed UDS on every spectral instrument we have tested, including the Sony ID7000™, the Cytek® Aurora, and the BD FACSDiscover™ S8.

### UDS can be quantified by unmixing reference data with multiple matrices

While UDS is often visible in fully-stained panel data, it can be difficult to attribute spread conclusively to UDS as opposed to other sources of noise like SSE from expression of other markers in the panel. In our analysis above, we showed that we can isolate the effects of UDS by instead evaluating a sample stained with a smaller backbone panel, not expected to have UDS issues, and unmixing it using either the backbone panel’s matrix or the larger panel’s matrix. Generalizing this technique, we propose a universal approach to quantifying UDS for a given panel: take raw reference data stained with only a subset of fluorochromes (or even unstained cells), unmix it with both the full panel matrix and a smaller reference matrix not expected to cause UDS, and compute and compare spread statistics (such as rSD) on the differently unmixed results (Figure 2A).

**Figure 2:**
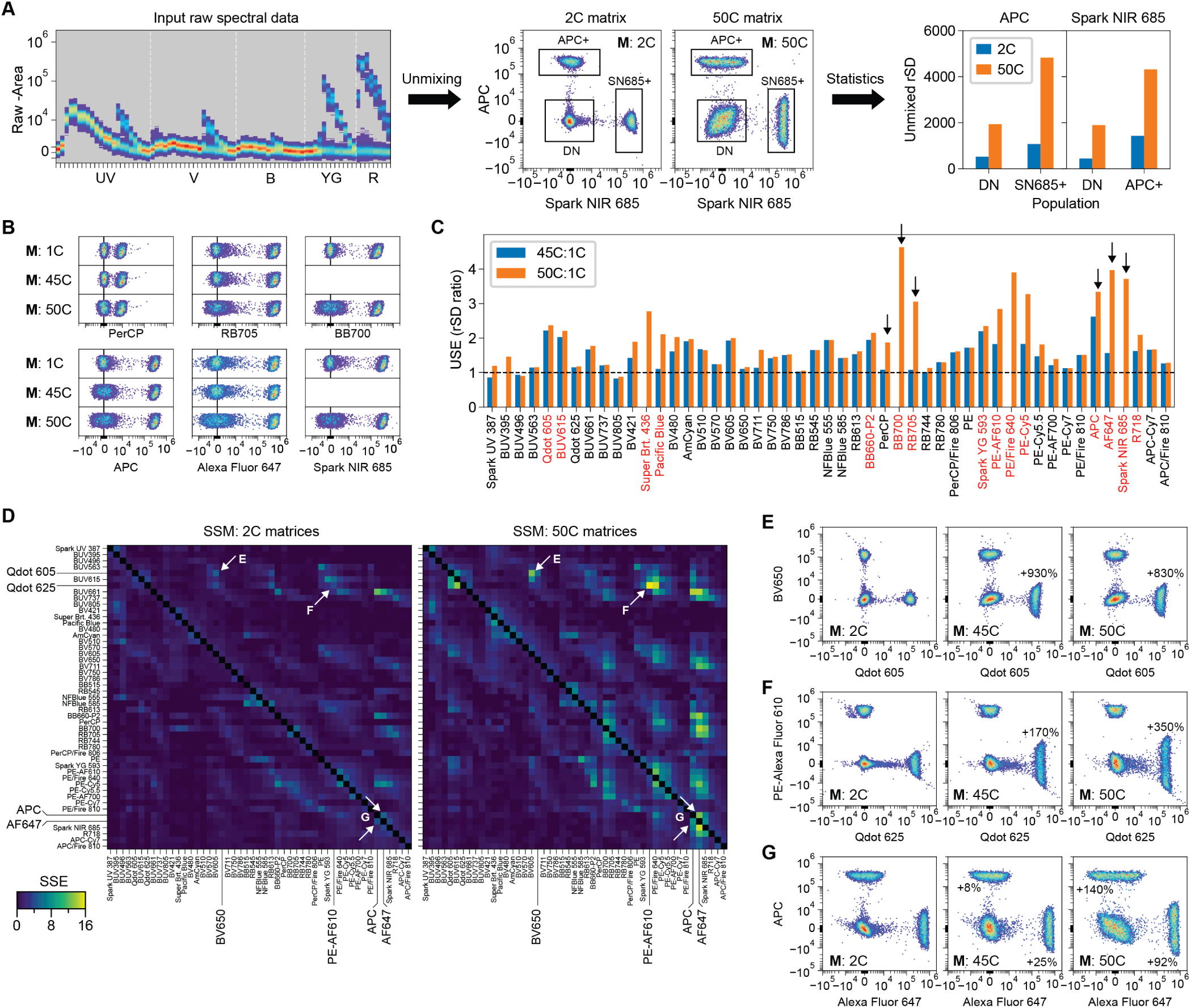
Quantifying UDS in OMIP-102. **(A)** Scheme: the same raw spectral data (left) is unmixed with multiple matrices (middle), one representing a reference condition with low UDS and one representing the panel of interest. Statistics related to spread, such as rSD, are calculated for specific populations under both unmixing conditions (right). **(B)** Visualization of USE for a subset of fluorochromes in the 45C and 50C versions of OMIP-102. For each fluorochrome, the same raw CD4 single-stain sample is unmixed using a single-color matrix containing that fluorochrome and AF (top subplot); a 45C matrix (middle subplot); and the full 50C matrix (bottom subplot). Vertical axis shows side scatter. **(C)** Calculation of USE for the fluorochromes in OMIP-102. An unstained sample is unmixed using each of the conditions described in (B), and the rSD of the unstained population for each unmixed fluorochrome is calculated. The ratio of the full panel (45C or 50C) unmixed rSD to the single-color unmixed rSD is reported. Arrows indicate examples shown in (B). Fluorochromes with >2-fold increase in rSD are highlighted in red. **(D)** Each entry in the “2C” spillover spreading matrix (SSM, left) is calculated using a matrix containing only the two fluorochromes for that entry, plus AF, as in (A). The “50C” SSM (right) is calculated using full-panel unmixing. White arrows indicate fluorochrome pairs with large differences in SSE between 2C unmixing and 50C unmixing. **(E)-(G)** Examples of fluorochrome pairs with high SSE in the 45C and 50C panels compared to 2C unmixing alone. Percent increase in SSE for single-positive populations is shown. All matrices also include AF as an unmixed parameter, with matrix size denoting the number of non-AF fluorochromes.

Specifically, we perform repeated unmixing of the same raw data of interest with two different spectral matrices. The first is a minimal “reference” matrix that accurately describes the sample of interest and with which UDS is expected to be minimal; for single-stain data, this could be a matrix containing only the present fluorochrome and AF. The second is the full-panel matrix including the full set of fluorochromes for the panel of interest, which may be suspected to induce UDS. The reference matrix should usually comprise a smaller subset of fluorochromes in the full-panel matrix. This process may be performed with data from a backbone panel, single-stained data, or even a single unstained recording. Regardless of the input sample, the matrices used should account for all fluorescence sources in that sample (including AF, if the full matrix includes AF) to ensure accurate unmixing. For example, a sample stained with a 20C backbone panel may be unmixed using a matrix containing those 20 colors, or using a matrix containing those 20C plus 31 additional colors, as in Figure 1A (we note that FMO controls are a well-known example of data being unmixed with a matrix containing strictly more fluorochromes than the sample). Obviously, the 51C stained sample should not be unmixed using the 20C backbone matrix, because it would fail to account for those 31 additional fluorochromes contributing signal in the data and would lead to unmixing errors that taint the subsequent statistical analysis.

Statistics calculated from this repeated unmixing approach can be used to derive metrics that quantify UDS, illustrated in Figure 2 using CD4 single-stain data for the fluorochromes in OMIP-102. We have previously described the “unmixing spreading error” metric (USE)^4^ for quantifying Hallmark 1. This metric is calculated by unmixing an unstained population with various single-color matrices (plus AF) for each color in the panel, and with the matrix for the full panel(s) of interest (Figure 2B, Supplementary Figure 2). The rSD of the unstained population can then be calculated for each unmixed parameter for the single-color and full-panel matrices, and USE is defined as the ratio of the full-panel unmixing rSD to the single-color unmixing rSD. USE can serve as a simple diagnostic metric for UDS (Figure 2C). This approach can be modified slightly to show the impact of UDS on single-color stain index directly (*e*.*g*., OMIP-102 Online Figure 4^4^). Any number of possible panel designs can be tested simultaneously by comparing reference single-color unmixing to two or more different panel designs, such as our comparison of spreading in OMIP-102’s 45C and 50C versions (Figure 2B-C). Similarly, Hallmark 2 (“tilted double-negatives”) can be quantified by computing the covariance and correlation matrices of an unmixed population for different matrices. The severity and direction of tilt for a given pair of unmixed parameters is captured by the magnitude and sign of the entry in the correlation matrix corresponding to that pair (Supplementary Figure 3).

The effect of UDS on spillover spreading described by Hallmark 3 can be quantified by comparing SSMs computed with different spectral matrices (Figure 2D-G). In this case, the reference unmixing condition for computing SSE for a given pair of fluorochromes is a 2C matrix containing those two fluorochromes, meaning that each entry in the left SSM in Figure 2D is computed with a different 2C matrix (plus AF). The SSM values can be inspected and compared directly as in Figure 2D, or the full-panel SSM can be reported as a percent change relative to the 2C SSM. This analysis reveals that, in some cases, UDS will lead to significant SSE between fluorochromes that show no measurable SSE unmixed as a pair (Figure 2E), while other cases show varying degrees of increase of SSE for pairs of fluorochromes that already had moderate-to-high SSE in isolation (Figure 2F-G). It is important to note that UDS simultaneously impacts the spread of both the positive and negative populations, meaning that metrics such as SSE, which subtract the variance fraction present in the negative population, may not reveal the full extent of UDS unless evaluated in tandem with other metrics such as USE.

We recommend calculating these empirical UDS metrics to check the outcome of unmixing both during the panel design process and after running a panel. They serve a similar role to FMOs by delineating the expected boundary between true positives and true negatives under actual measurement conditions (while FMOs are still needed to account for SSE from co-expression patterns). They also can inform panel design by indicating the effective resolution or stain index of a given fluorochrome in the context of the panel^4^ and predicting any expected areas of worse SSE that must be accounted for.

However, we acknowledge that single-stain data is often not available for all possible fluorochromes of interest early in the panel design process. To address that problem, we will discuss additional predictive metrics that can be calculated from publicly available fluorochrome data, such as spectral signatures on a given instrument.

### UDS tends to occur in spectral hotspots

In the example of OMIP-102, not all fluorochromes experience the same degree of USE, nor does the impact of UDS appear to be distributed randomly among fluorochromes (Figure 2C). Rather, fluorochromes with high unmixing spreading error tend to occur in clusters of spectrally neighboring fluorochromes (Figure 2C, red labels). When comparing UDS in the 45C and 50C versions of OMIP-102, we also observe that certain neighboring pairs of fluorochromes, such as APC and Alexa Fluor 647, have modest UDS in the 45C version but severe UDS in the 50C panel upon addition of a third or fourth fluorochrome in their spectral “neighborhood.” Because OMIP-102 was intentionally designed to avoid UDS as much as possible, these effects are somewhat subtle.

To build a deeper understanding of UDS, we next examined two iterations of a 40C panel differing by only one fluorochrome (Figure 3), where both panels have similar CN and distribution of fluorochrome similarities. The first panel, Panel 40C-A, overloads the “APC” spectral neighborhood (primarily red-excited fluorochromes emitting around 670nm) by including Vio Bright R667 along with APC and Spark NIR 685 (Figure 3A). Examination of unmixed CD4 single-stains for these fluorochromes using either 2C unmixing or full-panel unmixing shows substantial UDS affecting both the spread of the unstained population and the SSE of the positive populations (Figure 3B). By contrast, fluorochromes outside of the APC neighborhood (*e*.*g*., PE-Cy5.5 and BB700) are relatively unaffected (Figure 3B, bottom right). Quantification of USE confirms that UDS in Panel 40C-A is concentrated in the spectral neighborhood of APC (Figure 3C). We next evaluated Panel 40C-B, which differs from Panel 40C-A by only a single fluorochrome, replacing Vio Bright R667 with PerCP-Cy5.5 and thereby overloading the “BB700” neighborhood (primarily blue-excited fluorochromes emitting around 700nm) alongside BB700 and PE-Cy5.5 (Figure 3D). The fluorochromes in this neighborhood show severe UDS (Figure 3E), while the combination of APC and Spark NIR 685 from Panel 40C-A’s crowded neighborhood is less impacted (Figure 3E, bottom right). Most fluorochromes have similar USE in both panels, but the most severe UDS moves from the APC neighborhood to the BB700 neighborhood in the case of Panel 40C-B (Figure 3F). These example panels show the striking effect that adding or removing a single fluorochrome can have on both the severity and spectral location of UDS “hotspots” within a panel.

**Figure 3:**
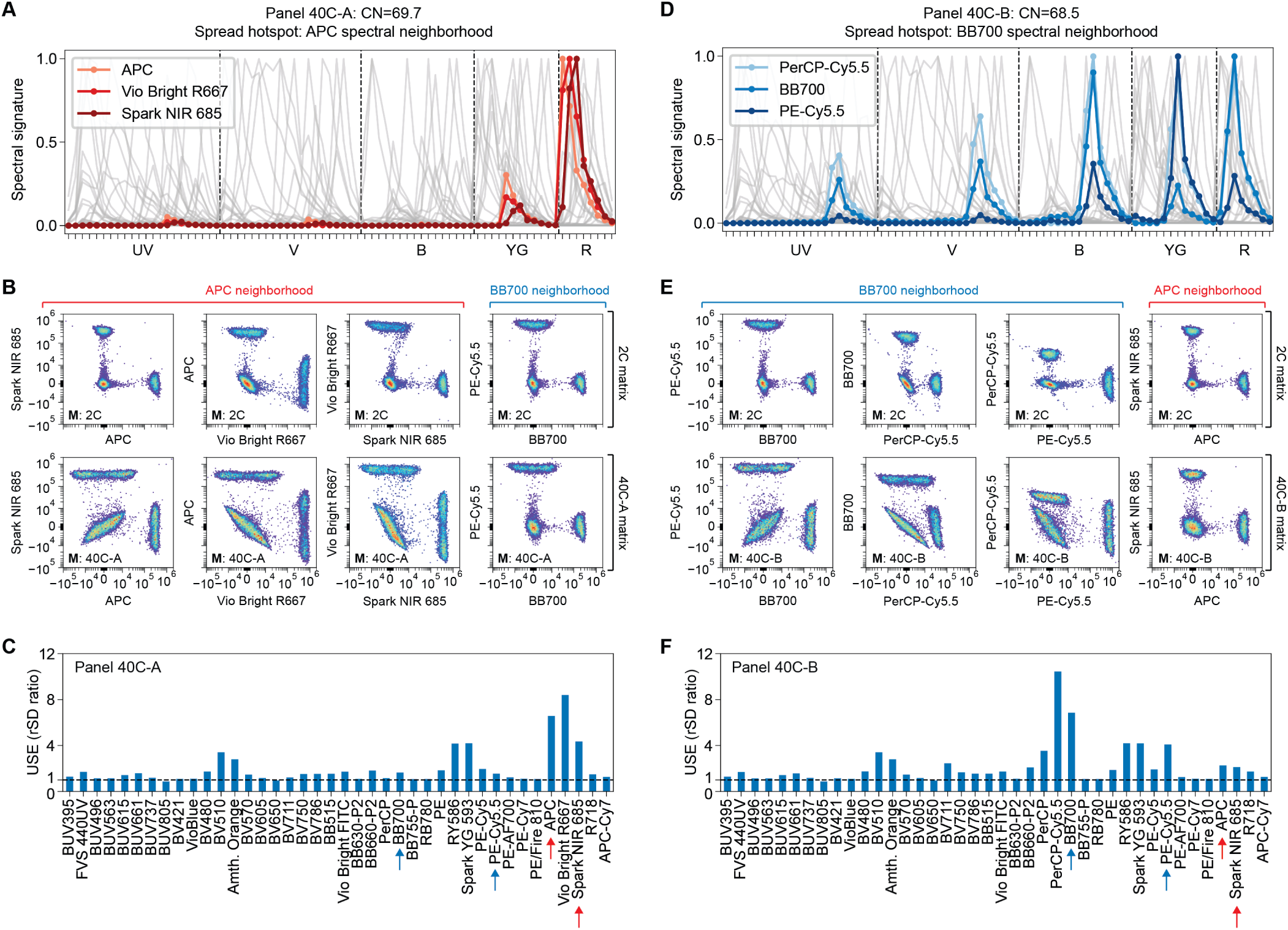
UDS is localized in panel-specific spectral “hotspots.” **(A)** Spectral signatures of Panel 40C-A, with fluorochromes in the “APC” spectral neighborhood highlighted in red. **(B)** Unmixed CD4 single-stain data for fluorochromes in the spectral neighborhood of APC show a large increase in spread when unmixed in the context of the full panel (bottom) compared to when unmixed as pairs (top). By contrast, a fluorochrome pair in the “BB700” spectral neighborhood in the same panel shows only a minor increase in spreading (bottom right). **(C)** USE calculated for the fluorochromes in Panel 40C-A. Larger spread is measured for fluorochromes in the APC neighborhood than in the BB700 neighborhood. **(D)** Spectral signatures for Panel 40C-B, a modified version of Panel 40C-A where Vio Bright R667 has been replaced with PerCP-Cy5.5. Spectra in the BB700 neighborhood are highlighted in blue. The condition number of the modified panel is essentially unchanged (68.5 vs 69.7). **(E)** Unmixed CD4 data as in (B) show that the most severe spreading “hotspot” now occurs in the BB700 spectral neighborhood for Panel 40C-B, while the APC neighborhood is less affected (right). **(F)** Quantification of USE confirms that the worst UDS occurs in the spectral neighborhood of BB700 for Panel 40C-B, not in the APC neighborhood as in Panel 40C-A. Arrows in (C) and (F) highlight the common subset of fluorochromes from both spectral neighborhoods that are present in both panels.

We next asked whether CS and CN can predict the presence, location, and severity of UDS hotspots in these panels. Figures 4A-B show that pairs of fluorochromes with high CS do not necessarily have comparable UDS in both panels. The CS of the APC and Spark NIR 685 pair is obviously unchanged between the two panels, but the presence of a third fluorochrome – Vio Bright R667 – causes a massive increase in spreading error in Panel 40C-A compared to Panel 40C-B. In Panel 40C-B, the fluorochrome pair with highest CS – BB515 and Vio Bright FITC – shows essentially no difference in UDS between 2C unmixing and full-panel unmixing (Figure 4B, inset). The introduction of PerCP-Cy5.5, whose CS with BB700 (0.91) is lower than the BB515 / Vio Bright FITC pair, seems to cause significant UDS. Meanwhile, a different fluorochrome pair with similarly high CS (RY586 and Spark YG 593, CS=0.91) has only moderate USE in both panels (Figure 3C, 3F). Fluorochromes with a high-similarity counterpart somewhere in the panel tend to have higher USE, but not always (Figure 4A-B, insets). Therefore, CS seems to be loosely predictive of a fluorochrome pair’s likelihood to be involved in UDS, but its limitation to pairwise relationships means that CS cannot reliably predict the actual fluorochrome combinations that cause UDS in a specific panel. This is problematic when attempting to design very large panels, where use of multiple high-similarity fluorochrome combinations is unavoidable.

**Figure 4:**
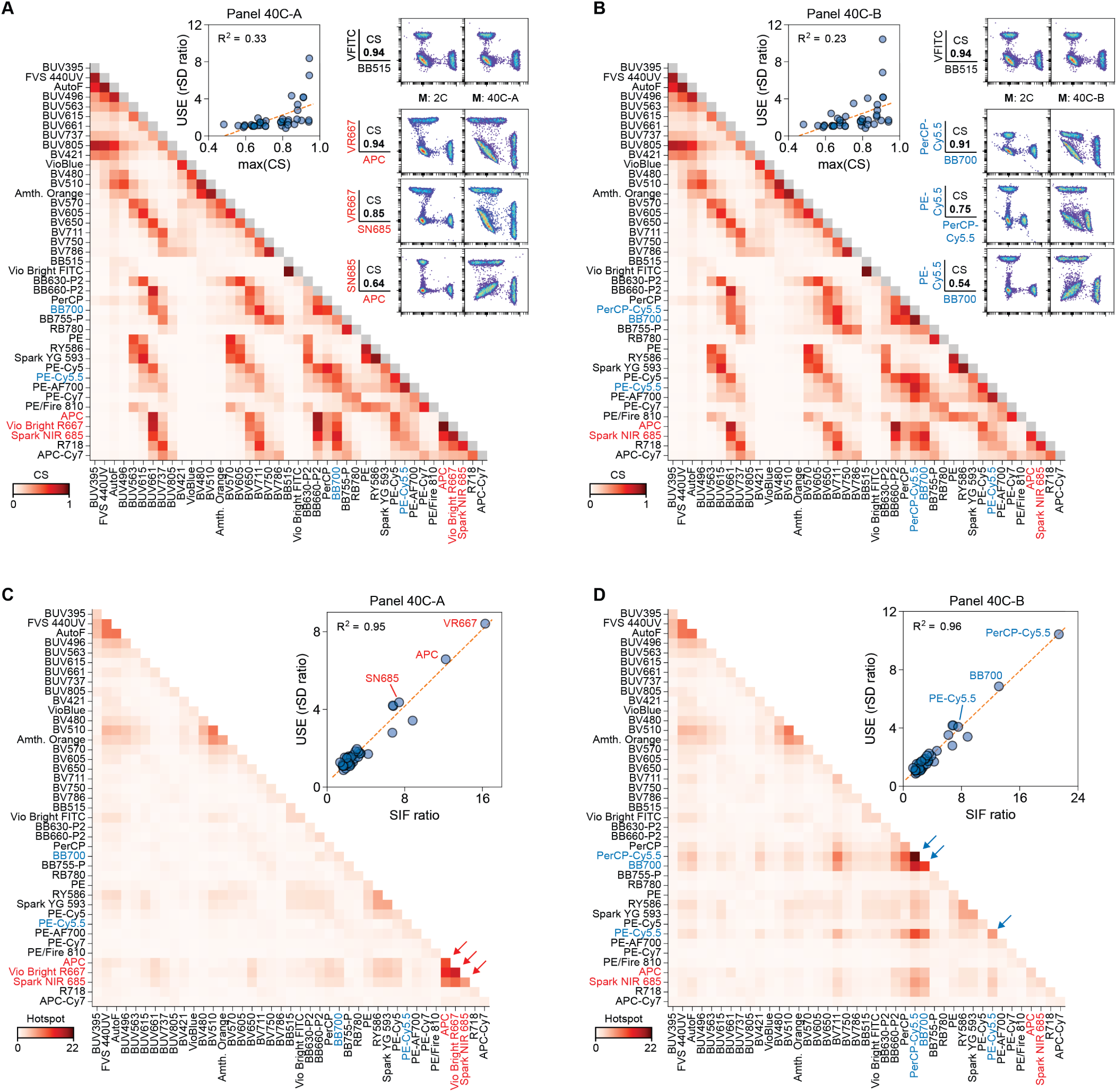
Predicting UDS with similarity and the Hotspot Matrix. **(A)-(B)** Cosine similarity (CS) matrices for Panels 40C-A **(A)** and 40C-B **(B)** do not clearly indicate the different locations of UDS hotspots in each panel. Measured USE is poorly predicted by each fluorochrome’s maximum CS (inset, center), with only some high-CS fluorochromes experiencing high USE. Example unmixed CD4 plots and CS values are shown for a high-similarity fluorochrome pair present in both panels (Vio Bright FITC and BB515) and the fluorochromes involved in each spreading “hotspot” from Figure 3 (inset, right) (“VFITC”: Vio Bright FITC; “VR667”: Vio Bright R667; “SN685”: Spark NIR 685). Of note, the high-similarity pair VFITC and BB515 shows no difference in UDS between 2C unmixing and 40C unmixing. **(C)-(D)** Hotspot Matrix analysis reveals panel-specific areas of UDS for Panels 40C-A **(C)** and 40C-B **(D)**. Diagonal values (Spreading Inflation Factors, SIFs) predict the degree of USE for each fluorochrome in the panel, while off-diagonal entries indicate which fluorochrome combinations are causing and affected by UDS. The APC and BB700 neighborhoods are uniquely indicated as hotspots (red and blue arrows) in Panels 40C-A and 40C-B, respectively, while several other less severe hotspots common to both panels are plainly visible. Insets show that predictive metrics derived from the Hotspot Matrix, such as the ratio of each fluorochrome’s panel-specific SIF to the SIF of that fluorochrome unmixed with a 1C+AF matrix alone, are strongly predictive of measured USE. R^2^ denotes the coefficient of determination for USE regressed against either maximum CS (A, B) or SIF ratio (C, D). “AutoF” denotes autofluorescence.

By contrast, condition number is predictive of UDS occurring somewhere in a panel, but gives no information about which fluorochromes are involved. Compared to Panels 40C-A and -B, a 39C subpanel containing neither Vio Bright R667 nor PerCP-Cy5.5 indeed has lower CN (39.0) and lower USE for fluorochromes involved in both the APC and BB700 hotspots (Supplementary Figure 4). Panels 40C-A and -B have nearly identical CN values (69.7 vs 68.5) that would be considered high relative to published 40C panels^20,21^. Unfortunately, CN alone gives no indication that UDS is localized in different parts of the spectrum across Panels 40C-A and -B. In summary, CN indicates if a panel is likely to have a UDS problem, but not which fluorochromes are involved; and CS may indicate whether or not a given two fluorochromes have the potential to experience UDS, but not to what extent they would be affected by UDS in a given panel. We will now use the fundamental mathematics of spectral unmixing to develop a new predictive metric that can diagnose and predict the severity of problematic combinations of fluorochromes in the context of a specific panel.

### The Hotspot Matrix mathematically predicts UDS based on spectral signatures alone

We can explain UDS mathematically by examining how uncertainty propagates through the spectral unmixing process. Fluorescence signals in a flow cytometer (either spectral or conventional) can be described with a linear mixture model^15,22^:

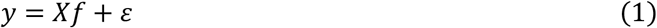

where *X* is the [*m* × *n*] spectral matrix containing *n* fluorochromes’ spectral signatures (columns of *X*) across *m* detectors (rows of *X*); *y* is the [*m* × 1] vector of measured raw detector signals for a given cell; *f* is the [*n* × 1] vector of fluorochrome abundances for that cell; and *ε* is a [*m* × 1] vector of random measurement noise.

Spectral unmixing (called “compensation” if *m* = *n*) is the process of solving Equation 1 for *f*, or, equivalently, finding an estimate of *f* that best explains the measured data *y* given the spectral matrix *X*. In the case of ordinary least squares (OLS) unmixing, the solution is given by:

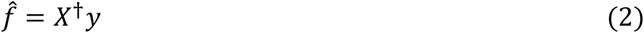

where the unmixed data vector 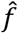 is the least-squares estimate of the true abundance values *f*, and *X*^†^ is the [*n* × *m*] Moore-Penrose pseudoinverse of *X*, where *X*^†^ = (*X*^*T*^*X*)^−1^*X*^*T*^.

Due to random measurement noise (*ε* in Equation 1), an individual cell’s measured spectrum will not be perfectly described by the spectral signatures in *X*, leading to some amount of error in the unmixed least-squares fit given by 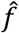 in Equation 2. Across many cells, this leads to statistical variance (“spread”) in the unmixed results around the average value. In the case of OLS unmixing, the relationship between raw measurement variance and unmixed variance can be expressed as:

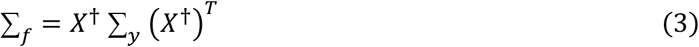

where ∑_*f*_ is the [*n* × *n*] covariance matrix of the unmixed fluorochrome abundances and ∑_*y*_ is the [*m* × *m*] covariance matrix of raw detector signals for a given set of cells (derivation in Supplementary Discussion). The diagonal values of the covariance matrices ∑_*f*_ and ∑_*y*_ are the squared standard deviations of each individual unmixed fluorochrome abundance or raw detector signal, respectively, while the off-diagonal terms describe covariances between pairs of fluorochromes or pairs of detectors. Equation 3 is exact, yielding results identical to what would be obtained by calculating statistics directly on unmixed data (Supplementary Figure 5).

We can draw two fundamental conclusions about UDS from the relation in Equation 3. First, UDS amplifies existing raw measurement noise, rather than introducing new noise; this is confirmed by simulations of noise-free raw data which reveal that subsequently unmixed data is also noise-free (Supplementary Figure 6). Second, Equation 3 confirms our empirical definition of UDS: the same raw data, with the same raw covariance matrix, can result in different unmixed variance if a different spectral matrix is used for unmixing (Supplementary Figure 7, further explanation in Supplementary Discussion). These observations lie at the heart of UDS, providing a clear mathematical explanation for its seemingly unintuitive empirical hallmarks (Supplementary Figure 8).

Based on these insights, we want to derive a new predictive metric for UDS that captures the impact of the spectral matrix *X* alone, independent of the specific input raw data described by ∑_*y*_. To do this, we assume a simplified form of ∑_*y*_ in which noise is uncorrelated across detectors and every detector has the same average variance *σ*^2^ (these simplifications, which OLS also assumes, are not generally true for flow cytometry data, but are reasonable enough for predictive purposes). Then ∑ = *σ*^2^*I*, where *I* is the [*m* × *m*] identity matrix, and Equation 3 reduces to the following (derivation in Supplementary Materials):

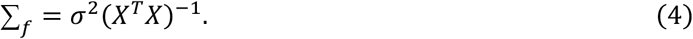

Equation 4 reveals that unmixed covariance is proportional to the matrix (*X*^*T*^*X*)^−1^. We note that if the columns of *X* are normalized to have unit length, then *X*^*T*^*X* is simply the “similarity matrix” containing each fluorochrome pair’s CS (equivalent to the matrix of Pearson correlation coefficients if the columns of *X* are also centered to have a mean of zero); the matrix (*X*^*T*^*X*)^−1^ is therefore the inverse of the similarity matrix. The magnitude of UDS (in “standard deviation” data units, as opposed to variance’s squared data units) is thus proportional to:

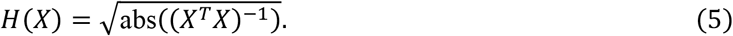

We have termed *H*(*X*) the “Hotspot Matrix” for its ability to predict empirical UDS hotspots in a given panel of fluorochromes when visualized as a heatmap. Figures 4C-D show such a visualization for Panels 40C-A and 40C-B, in which the fluorochromes involved in the APC and BB700 “hotspots” are immediately obvious from visual inspection of each panel’s corresponding Hotspot Matrix. More intense hotspots correspond to more severe regions of UDS in each panel, while less intense hotspots reveal moderately problematic fluorochrome combinations.

The diagonal values of *H*(*X*) indicate the factor by which each fluorochrome’s unmixed standard deviation is expected to increase when unmixed with the spectral matrix *X*, relative to being unmixed with a matrix containing that fluorochrome alone. We term these fluorochrome-specific diagonal values “Spreading Inflation Factors” or “SIFs,” based on a mathematically identical metric described in the regression literature^23,24^ called the “Standard Error Inflation Factor” (square root of the more commonly known “Variance Inflation Factor”^25,26^). The magnitude of each fluorochrome’s SIF indicates the extent to which that fluorochrome’s unmixed resolution is degraded by UDS (Supplementary Figure 9). In practice, we have found the following rough guidelines to be helpful: SIFs below 3 are generally of minimal concern; SIFs between 3 and 8 indicate moderate UDS that should be managed in the panel design process; and SIFs exceeding 8 indicate severe UDS. To provide a dimensionless metric with similar interpretation to USE, we computed the “SIF ratio,” the ratio of full-panel unmixing SIFs to single-color unmixing SIFs (Supplementary Figure 9). Despite being computed solely from spectral signatures, SIF ratios are remarkably strong predictors of measured USE (Figure 4C-D, insets), showing the utility of the Hotspot Matrix for predictive analysis during the panel design process.

While large diagonal entries in *H*(*X*) identify individual fluorochromes involved in UDS, large off-diagonal entries reveal the specific combinations of fluorochromes that are implicated in UDS (Figure 4C-D). These off-diagonal entries are proportional to the square root of the magnitude of covariance between pairs of unmixed parameters, and therefore indicate the severity of the “tilted double-negative” effect described by Hallmark 2. In fact, both the direction and severity of tilt can be predicted from the signs and magnitudes of entries in (*X*^*T*^*X*)^−1^ (Supplementary Figure 10). When multiple hotspots are present, noticeable off-diagonal entries may be visible at the intersection of non-overlapping spectra (*e*.*g*., PerCP-Cy5.5 and AF in Figure 4D). In these cases, the actual fluorochrome combinations causing UDS will be indicated by larger off-diagonal values elsewhere for the same fluorochromes (*e*.*g*., PerCP-Cy5.5 with BB700, and AF with Fixable Viability Stain 440UV in Figure 4D). To summarize, UDS in a panel can be detected with the Hotspot Matrix following a two-step procedure: (1) large diagonal values identify which fluorochromes are affected by UDS, and to what extent; and (2) large off-diagonal values identify the combinations of fluorochromes that are causing UDS in the first place.

The Hotspot Matrix provides a predictive tool for diagnosing and quantifying the impact of UDS on a given panel, while simultaneously identifying the problematic combinations of fluorochromes that lead to UDS. To demonstrate its general applicability, we have applied Hotspot Matrix analysis to predict and identify UDS across a range of spectral cytometers and panels, including OMIP-102^4^, OMIP-069^14^, and OMIP-109^27^ (Supplementary Figures 11-15). In the case of OMIP-069, we note that the inclusion or exclusion of AF as an unmixed parameter can impact the accuracy of USE prediction with the Hotspot Matrix for fluorochromes whose spectra substantially overlap that of AF (Supplementary Figure 15).

Figure 5 illustrates how Hotspot Matrix analysis can be used to iteratively identify and replace problematic fluorochrome combinations as part of the panel design process. We repaired Panel 40C-A following a simple procedure: remove a fluorochrome involved in the worst hotspot, and replace it with a different fluorochrome that does not create a new hotspot or worsen an existing hotspot. After just four substitutions, CN was reduced from 69.7 to 15.7 while USE was reduced to below 2.5 for all fluorochromes (Figure 5E). Subsequently adding an additional 5 fluorochromes to the panel (Figure 5F), again guided by Hotspot Matrix analysis, was possible with virtually no impact to measured USE and only a modest increase in CN (19.6). This example highlights how using UDS-related predictive metrics before starting any wet-lab work can avoid or reduce costly trial-and-error experiments during the panel design process.

**Figure 5:**
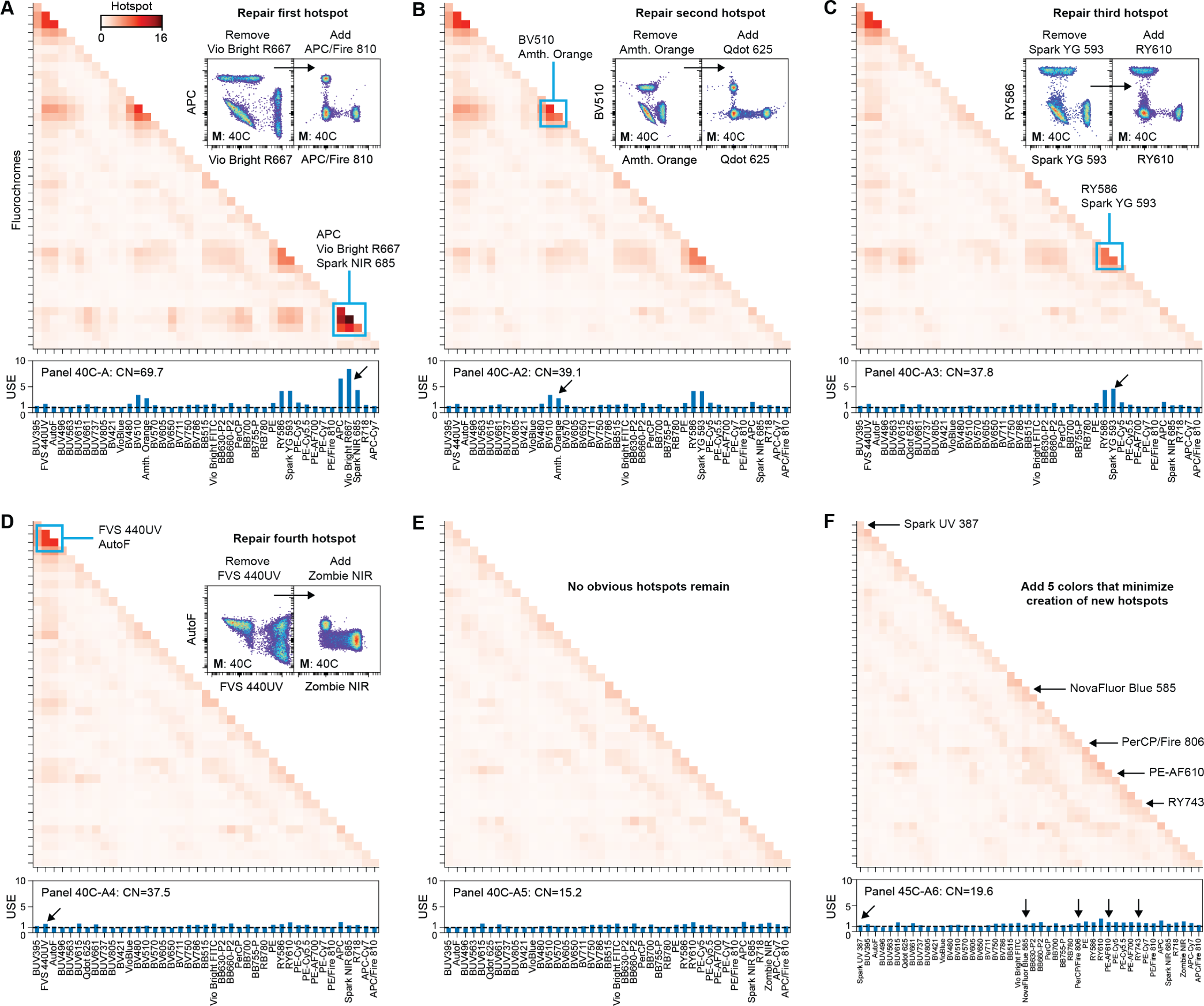
Iterative repair and in-silico expansion of a 40C panel using Hotspot analysis. **(A)** We begin by identifying the most severe hotspot in Panel 40C-A, confirmed with measured USE (bottom). By replacing one fluor in the hotspot (Vio Bright R667) with a new fluor that does not induce a hotspot (APC/Fire 810), the original hotspot is removed and the panel CN improves as shown in (B). **(B)-(D)** We repair the panel iteratively by identifying a severe remaining hotspot, removing one fluorochrome in that hotspot, and replacing it with an appropriate substitute that does not induce a new hotspot. **(E)** Eventually, all obvious hotspots are removed, resulting in a panel with low CN and low USE. **(F)** The same strategy can be used to expand the panel, here from 40C to 45C, by iteratively searching for fluorochromes that do not induce a new hotspot when added. Using this strategy, we expanded from 40C to 45C while increasing CN from 15.2 to only 19.5, with minimal increase in USE.

### Spectral collinearity is the underlying cause of UDS

We conclude with an explanation of the underlying mathematical cause of UDS in terms of the statistical concepts of “variance inflation” and “collinearity.” Variance inflation occurs in multiple regression when highly correlated predictor variables lead to increased uncertainty in regression parameter estimates^18,25^. Descriptions of variance inflation in the statistics literature are remarkably similar to the empirical hallmarks of UDS, such as higher-than-expected variance in regression results when using correlated predictor variables^18^ (Hallmark 1) and high covariance between regression coefficients leading to visible “tilt” in bivariate plots^28,29^ (Hallmark 2). Variance inflation is known to be caused by predictor variables with high collinearity, a term describing the degree of intercorrelation among a set of variables or vectors^18^.

Approaches for diagnosing collinearity and variance inflation can be applied to the problem of UDS by framing spectral unmixing as a multiple regression problem. In this interpretation, unmixing is the process of fitting a linear regression model that explains a cell’s measured raw signal *y* (the dependent “response variable” or “regressand”) across *m* detectors (“observations”) in terms of *n* fluorochrome spectra (independent “predictor/explanatory variables” or “regressors”, summarized in matrix *X*). Each individual cell’s estimated fluorochrome abundances 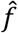 are therefore given by the regression “parameters” or “coefficients” that best fit the model to that cell’s raw signal. While this statistical interpretation of unmixing is mathematically identical to the linear mixture model we described previously, it allows a new understanding of UDS as a process of variance inflation arising from collinear combinations of spectra. Just like practitioners of multiple regression work to prevent problematic variance inflation by avoiding excessive correlation among predictor variables^18,25,30^, the spectral cytometrist seeks to avoid or mitigate UDS arising from collinearity among spectra in their panel.

Collinearity not only describes the obvious case of two overly similar spectra (whose vectors are nearly “co-linear,” or parallel, with a CS approaching 1 – see Supplementary Figure 1), but also generalizes the pairwise concepts of correlation and similarity to combinations of three or more spectra^18,30^. Mathematically, collinear combinations of spectra are nearly “linearly dependent” on one another, such that one spectrum can almost be expressed as a linear combination (weighted sum) of one or more other spectra, leading to an ill-conditioned system. The “APC” and “BB700” spectral hotspots identified in Panels 40C-A and 40C-B are examples of collinear combinations of three spectra, which can be visualized geometrically as three vectors which nearly fall in the same two-dimensional plane^30^ (Supplementary Figure 16).

The literature on variance inflation echoes our earlier observations that CN, as a single aggregate metric, cannot identify the specific sources of collinearity in a panel^24,30,31^, while CS, as a pairwise metric, cannot identify collinearity arising from combinations of more than two spectra^30–32^. Metrics based on the inverse correlation matrix of predictor variables (from which the Hotspot Matrix is derived), like VIFs and SIFs, have been used extensively in the regression literature to diagnose collinearity^23–25,30,31^. We note that our suggested interpretation of off-diagonal Hotspot Matrix values has been previously described and supported^31^. Additional diagnostic techniques based on matrix decomposition, such as the Variance Decomposition Proportion (VDP)^18,30,33^, can be combined with Hotspot Matrix analysis to provide unambiguous identification of collinear spectra, even when multiple overlapping collinearities are present (Supplementary Figure 17).

## DISCUSSION

UDS can have a profound impact on the final resolution and quality of unmixed data in a spectral panel. Here we have shown how UDS can be explained, quantified, predicted, and ultimately mitigated as part of the panel design process. The Hotspot Matrix both predicts the magnitude of UDS in a spectral panel and identifies the specific collinear combinations of fluorochromes that cause UDS, providing a platform-agnostic tool to design panels that avoid problematic levels of UDS. Because it depends only on fluorochrome spectral signatures, the Hotspot Matrix can be employed for *in silico* panel design without the need for measured data from an individual cytometer, especially since the use of standardized instrument settings and optical configurations allows the use of “representative” pre-recorded spectral signature data for a specific instrument platform. In the panel design workflow, the Hotspot Matrix is primarily useful when evaluating compatible combinations of colors and assessing the relative impact of UDS on marker resolution, but it does not replace existing panel-design considerations such as matching fluorochrome brightness and antigen density, or accounting for co-expression when managing SSE (summary of spectral panel design metrics and terminology in Table 1). We also recommend quantifying UDS in measured data as a quality-control step when evaluating unmixing performance in a panel, alongside existing QC metrics and staining controls.

**Table 1:** Summary of published spectral panel design metrics related to unmixing-dependent spreading (UDS). The “Fluorochrome scope” column indicates whether the metric refers to single fluorochromes, pairs of fluorochromes, or full panels of fluorochromes. The “Panel-specific” column indicates whether or not the value of the metric depends on the total panel of fluorochromes in the spectral matrix. The “Data required” column indicates what input data is needed in order to calculate the metric.

| Metric / matrix name | Alternative names | Fluorochrome scope | Panel-specific | Data required | Cytometry description | Mathematical description | References |
| --- | --- | --- | --- | --- | --- | --- | --- |
| Cosine similarity (CS) / similarity matrix | “similarity index,” “similarity scores,” “similarity,” “uncentered correlation matrix” | Pairwise | No | Spectral signatures only | Metric describing the degree of spectral overlap or similarity between two fluorochromes, based on their normalized spectral signatures. Spectra with CS=0 have no overlap, while spectra with CS=1 are spectrally identical. | Cosine of the angle between two fluorochromes’ spectral signatures, calculated as the dot product of their signatures normalized to unit length. Equivalent to an uncentered correlation coefficient. Spans [0,1] for non-negative spectral signatures. | Kotu 2019<sup>17</sup>, Park 2020<sup>14</sup> |
| Condition number (CN) | “complexity index,” “complexity score,” “complexity” | Full panel | Yes | Spectral signatures only | Single metric describing the overall difficulty of unmixing a specific combination of fluorochromes. Smaller numbers are better. | Describes the numerical sensitivity of the unmixing problem, *i.e.*, how much the unmixed result will change for a small perturbation in the input. Calculated as the ratio of the maximum to the minimum singular value of the spectral matrix. Spans [1, ∞]. | Belsley 1980<sup>18</sup>, Park 2020<sup>14</sup> |
| Hotspot Matrix | - | Pairwise and single-color | Yes | Spectral signatures only | Matrix describing the extent to which different fluorochromes are affected by UDS, and which combinations cause UDS. | Inverse of similarity matrix, or product of spectral matrix pseudoinverse with its transpose matrix. Diagonal spans [1, ∞]. Off-diagonal spans [0, ∞]. | This work |
| Spreading Inflation Factor (SIF) | “collinearity index”<sup>24</sup>, “standard-error inflation factor”<sup>23</sup> | Single-color | Yes | Spectral signatures only | Predicted increase in spread (rSD) of an unmixed fluorochrome in the context of a specific panel’s matrix. | Diagonal of hotspot matrix; square root of Variance Inflation Factor. Spans [1, ∞]. | This work, Stewart 1987<sup>24</sup>, Fox 1992<sup>23</sup> |
| Spreading error (SE) | - | Single-color | Yes | Unmixed data | General term describing measured variance, or “spread,” in unmixed or compensated data. | rSD or related metric describing an unmixed parameter corresponding to a specific population. | Roederer 2001<sup>35</sup> |
| Unmixing spreading error (USE) | - | Single-color | Yes | Unmixed data | An increase in spread of an unmixed parameter across all cells when unmixed with a full-panel matrix, regardless of marker expression. | Ratio of an unstained population’s rSD unmixed with a full panel matrix to its rSD unmixed with a matrix containing just the single color and AF. | Konecny 2024<sup>4</sup>, this work |
| Spillover spreading error (SSE) / Spillover spreading matrix (SSM) | - | Pairwise | Yes | Unmixed data | Increase in the spread of one fluorochrome’s unmixed signal due to emission from a second fluorochrome that spectrally overlaps the first, normalized to the intensity of the second fluorochrome “causing” spread. | Square root of the difference in unmixed variance for one fluorochrome between a positive and a negative population for a second fluorochrome, divided by the difference in MFI of the second fluorochrome between both populations. | Nguyen 2013<sup>8</sup> |
| Total spreading error (TSE) / Total spreading matrix (TSM) | - | Pairwise | Yes | Unmixed data | Same as SSE, but not normalized to the intensity of the fluorochrome “causing” spread. | Same as SSE, but not normalized to the delta-MFI of the positive fluorochrome. | Corselli 2023<sup>36</sup> |

We note several limitations and caveats to consider when employing Hotspot Matrix analysis. First, because it relies on the simplifying assumptions of the OLS noise model, the Hotspot Matrix is not a perfect quantitative predictor of absolute measured UDS (as reflected by moderate dispersion in Figure 4D, and the fact that SIF ratios can overestimate measured USE by roughly 2-fold). For this reason, rather than attempting exact quantitative predictions, we recommend using the Hotspot Matrix to check for the presence or absence of severe UDS, and to assess the relative impact of UDS on different fluorochrome combinations. We have provided approximate SIF thresholds for identifying problematic collinearity based on our experience, but ultimately the reader must determine what degree of UDS is acceptable for a given fluorochrome in the context of their panel (similar subjectivity is noted in the regression literature when attempting to define absolute thresholds for problematic collinearity^30^). Although based on the OLS noise model, the Hotspot Matrix still provides useful relative indications of collinearity for data unmixed via non-OLS algorithms (Supplementary Figure 18). Finally, because a fluorochrome’s spectral signature in a given experiment will depend on the instrument settings used, Hotspot Matrix values (along with CS and CN) may vary when using detector gains that deviate significantly from default settings.

As the menu of fluorochromes available for flow cytometry continues its rapid expansion, tools for predicting UDS will enable the increasingly important task of identifying a good palette of fluorochromes for the given plexity of a panel. Our workflow allows this initial step of fluorochrome selection to be performed *in silico* in a platform-agnostic fashion by leveraging publicly available spectral signature datasets. Only after identifying fluorochrome combinations that produce minimal UDS for a given panel size does the experimenter need to move to experimental reagent and panel testing to ensure that the sought-after resolution for all markers in the panel can be achieved. Moving forward, UDS prediction tools like the Hotspot Matrix have potential utility throughout the spectral cytometry workflow, such as the identification of optimal subsets of autofluorescence spectra for unmixing in heterogeneous samples^34^.

## Acknowledgments

We would like to thank Shirley Shi, Tri Le, Aaron Middlebrook, Chip Lomas, Jessica Stokes, and Emilie Jalbert (BD Biosciences) for assistance with sample preparation and data collection. We also thank Gisele Baracho (BD Biosciences) for panel design assistance. We thank Laura Ferrer (BD Biosciences) for review and writing assistance. We thank Aaron Tyznik (BD Biosciences) and Martin Prlic (Fred Hutchinson Cancer Center) for organizational and administrative support. This work was supported in part by NIH grants R01 AI123323 and R56 DE032009 (to Martin Prlic). P.L.M. is an ISAC International Innovator. F.M. is a former ISAC Marylou Ingram Scholar.

## Author Contributions

Peter L. Mage: Conceptualization; data curation; mathematical derivations; formal analysis; methodology; software; validation; visualization; writing - original draft; writing - review and editing.

Andrew J. Konecny: Conceptualization; data curation; writing - original draft; writing - review and editing. Florian Mair: Conceptualization; writing - original draft; writing - review and editing.

