## Supplementary Materials for "Measurement and prediction of unmixing-dependent spreading in spectral flow cytometry panels"

### SUPPLEMENTARY METHODS

#### Summary of datasets

Supplementary Table S1 summarizes which data set and what fluorochromes were used for illustration in each figure. Original data from OMIP-102 was re-analyzed to demonstrate novel concepts in the current manuscript.

#### Staining procedure for full stain panels and single stain controls

Previously unpublished 20C and 51C panel data from the development of OMIP-102 (Figure 1A) were prepared following an identical procedure previously described in OMIP-102[1]. When necessary, intracellular staining (Cytofix/Cytoperm™ Fixation/Permeabilization Kit, BD 554714) or intranuclear staining (eBioscience™ Foxp3 / Transcription Factor Staining Buffer Set, Thermo Fisher 00-5523-00) was additionally performed according to the manufacturer's recommendations. Cryopreserved peripheral blood mononuclear cells (PBMCs) were obtained from whole blood or leukopak purchased from Bloodworks Northwest (https://www.bloodworksnw.org/, Washington USA). Different donors were used in the generation of full stain panels and single stain controls.

Previously unpublished 25C and 40C panel data (Figures 1B, D-F) and anti-CD4 single-stain data were prepared using fresh PBMCs isolated from whole blood from healthy donors. Blood was collected in EDTA anticoagulant-containing tubes (BD) and diluted with an equal volume of PBS (Corning Inc.). Diluted blood was loaded on top of 15-mL density gradient medium Ficoll™ (Cytiva) in a SepMate™ PBMC Isolation Tube (STEMCELL Technologies). Centrifugation was performed at 1200 × *g* for 10 min with brake on at room temperature. After collection of the buffy coat, cells were washed with PBS twice and resuspended at 10 million cells/mL. Anti-CD4 staining was performed at room temperature for 30 minutes in the dark, followed by two washes in PBS. Staining was performed according to manufacturer recommendations, including recommended additives to prevent fluorochrome interactions and aggregation. All experiments using human whole blood from healthy donors were done in compliance with the criteria of the Environment Health and Safety office in BD Biosciences.

#### Cytometers used and acquisition settings

New and reanalyzed published data were derived from four full spectrum cytometers: a 5-laser instrument with a total of 48 fluorescent detectors (commercially available from BD Biosciences as the BD FACSymphony™ A5 SE); a 5-laser instrument with a total of 78 fluorescent detectors (commercially available from BD Biosciences as the BD FACSDiscover™ S8); a 5-laser instrument with a total of 64 fluorescent detectors (commercially available from Cytek Biosciences as the Cytek® Aurora); and a 7-laser instrument with a total of 184 fluorescent detectors (commercially available from Sony Biotechnology as the Sony ID7000™).

Detector gains were set as follows for each cytometer. For the Cytek® Aurora, the default "Cytek® Assay Settings" as chosen by the manufacturer were used. For the Sony ID7000™, detector gains were determined via a voltration-like method as described in OMIP-102 with the 808 nm laser turned off, therefore utilizing 182 out of the 184 fluorescent detectors. For the BD FACSymphony™ A5 SE, detector voltages were selected so that the rSD of unstained lymphocytes in each channel would equal approximately 3× the rSD of electronic noise determined during cytometer setup.

For the BD FACSDiscover™ S8, default APD gains targeting pre-defined MFIs for a hard-dyed setup bead were used as chosen by the manufacturer. Anti-CD4 single stain data on the S8 was acquired on one of three different instruments with matched optical configurations, with default gains matching the same target MFIs for a hard-dyed setup bead. We note that the manufacturer updated the default target MFIs in BD FACSChorus 5.2 (May 2024), resulting in some anti-CD4 single-stain recordings with higher gains than others in this work. The updated target MFIs included a fixed global increase in gain across all detectors, preserving the shape of spectral signatures before and after the settings update and allowing combined analysis of spectral signatures from data acquired at either set of gains.

#### Analysis and gating

Unless otherwise noted, analysis was performed on lymphocytes defined via gating on forward- and side-scatter parameters.

### SUPPLEMENTARY DISCUSSION

#### Dependence of unmixed variance on pseudoinverse matrix

Following Equation 1 (main text), the signal $y_i$ on detector $i$ is a linear combination, or weighted sum, of each fluorochrome's relative contribution to that detector ($x_{ij}$ describes the signal from fluorochrome $j$ into detector $i$, relative to fluorochrome $j$'s maximum detector), weighted by each fluorochrome's abundance $f_j$:

$$y_i = \sum_{j}^{n} x_{ij} f_j = x_{i0} f_0 + x_{i1} f_1 + \cdots + x_{in} f_n \qquad \text{(SI-1)}$$

We can express the OLS-unmixed abundance of a single fluorochrome (Equation 2, main text) as follows:

$$\hat{f}_j = \sum_{i}^{m} x_{ji}^{\dagger} y_i = x_{j0}^{\dagger} y_0 + x_{j1}^{\dagger} y_1 + \cdots + x_{jm}^{\dagger} y_m \qquad \text{(SI-2)}$$

where $x_{ji}^{\dagger}$ is the entry of the pseudoinverse matrix $X^{\dagger}$ corresponding to fluorochrome $j$ and detector $i$. Just like a fluorochrome's spectrum (column of $X$) describes how its signal maps to detectors (Equation SI-1), a fluorochrome's "inverse spectrum" (row of $X^{\dagger}$) describes how raw detector signals map back to its unmixed abundance (Equation SI-2). A fluorochrome's spectrum in $X$ is a fixed property of that fluorochrome on a given instrument and does not depend on the other fluorochromes in $X$, but a fluorochrome's inverse spectrum in $X^{\dagger}$ will vary depending on the overall set of fluorochromes in the matrix used for unmixing (Supplementary Figure 7). Conceptually, this occurs because the unmixing estimation problem is different depending on the other fluorochromes in the matrix.

It is instructive to break down the matrix expression in Equation 3 (main text) to see how a single fluorochrome's unmixed variance depends on both raw detector covariances and its inverse spectrum. If we make the common assumption that detector noise is uncorrelated and therefore $\sum_y$ has no nonzero off-diagonal elements, we have:

$$\sigma_{f_j}^2 = \sum_{i}^{m} \left(x_{ji}^{\dagger}\right)^2 \sigma_{y_i}^2 \qquad \text{(SI-3)}$$

where $\sigma_{f_j}^2$ is the variance of fluorochrome $j$ and $\sigma_{y_i}^2$ is the variance of detector $i$. A given fluorochrome's unmixed variance is therefore the sum of each raw detector's variance, weighted by the squared magnitude of the fluorochrome's inverse spectrum value for that detector. An inverse spectrum with large-magnitude values (positive or negative) will result in more significant amplification of raw noise (compare hotspot fluorochromes in Figure 3C and 3F with inverse spectra in Supplementary Figure 7). Just as Equation SI-2 describes how a fluorochrome's inverse spectrum maps signal from detectors to its unmixed abundance, Equation SI-3 shows that the inverse spectrum also maps noise from raw detectors to unmixed abundance. Equation SI-3 also explains Hallmark 3 of UDS, wherein SSE can be selectively increased in certain panels if the magnitudes of inverse spectra values for a "receiving" fluorochrome are increased for detectors with high emission from a "SSE-causing" fluorochrome (Supplementary Figure 8).

#### Estimator covariance derivations

Because $\hat{f}$ is an unbiased estimator of $f$, the expected value of $\hat{f}$ is simply $E[\hat{f}] = f$. Then the variance of $\hat{f}$ is given by

$$
\begin{aligned}
\Sigma_f = V[\hat{f}] &= E\left[\left(\hat{f} - E[\hat{f}]\right)^2\right] \\
&= E\left[\left(\hat{f} - f\right)^2\right] \\
&= E\left[\left(\hat{f} - f\right)\left(\hat{f} - f\right)^T\right]
\end{aligned}
$$

The difference between estimated abundance $\hat{f}$ and the true abundance $f$ is given by:

$$
\begin{aligned}
\hat{f} - f &= X^\dagger y - f \\
&= X^\dagger (Xf + \varepsilon) - f \\
&= f + X^\dagger \varepsilon - f \\
&= X^\dagger \varepsilon
\end{aligned}
$$

Therefore

$$
\begin{aligned}
V[\hat{f}] &= E\left[X^\dagger \varepsilon \left(X^\dagger \varepsilon\right)^T\right] \\
&= X^\dagger E[\varepsilon \varepsilon^T] \left(X^\dagger\right)^T
\end{aligned}
$$

We can prove that $E[\varepsilon\varepsilon^T]$ is simply the variance of $y$, recalling that for the OLS solution, $E[\varepsilon] = 0$:

$$
\begin{aligned}
\Sigma_y = V[y] &= E[(y - E[y])^2] \\
&= E[(Xf + \varepsilon - E[Xf + \varepsilon])^2] \\
&= E[(Xf + \varepsilon - Xf - 0)^2] \\
&= E[(\varepsilon)^2] \\
&= E[\varepsilon \varepsilon^T]
\end{aligned}
$$

Therefore, $\Sigma_f = X^\dagger \Sigma_y \left(X^\dagger\right)^T$, confirming Equation 3 in the main text.

For OLS, we have $\Sigma_y = \sigma^2 I$. Then:

$$\begin{aligned}\Sigma_f &= X^\dagger \sigma^2 I \left(X^\dagger\right)^T \\ &= \sigma^2 X^\dagger \left(X^\dagger\right)^T \\ &= \sigma^2 (X^T X)^{-1}\end{aligned}$$

This confirms Equation 4 in the main text.

### SUPPLEMENTARY FIGURES

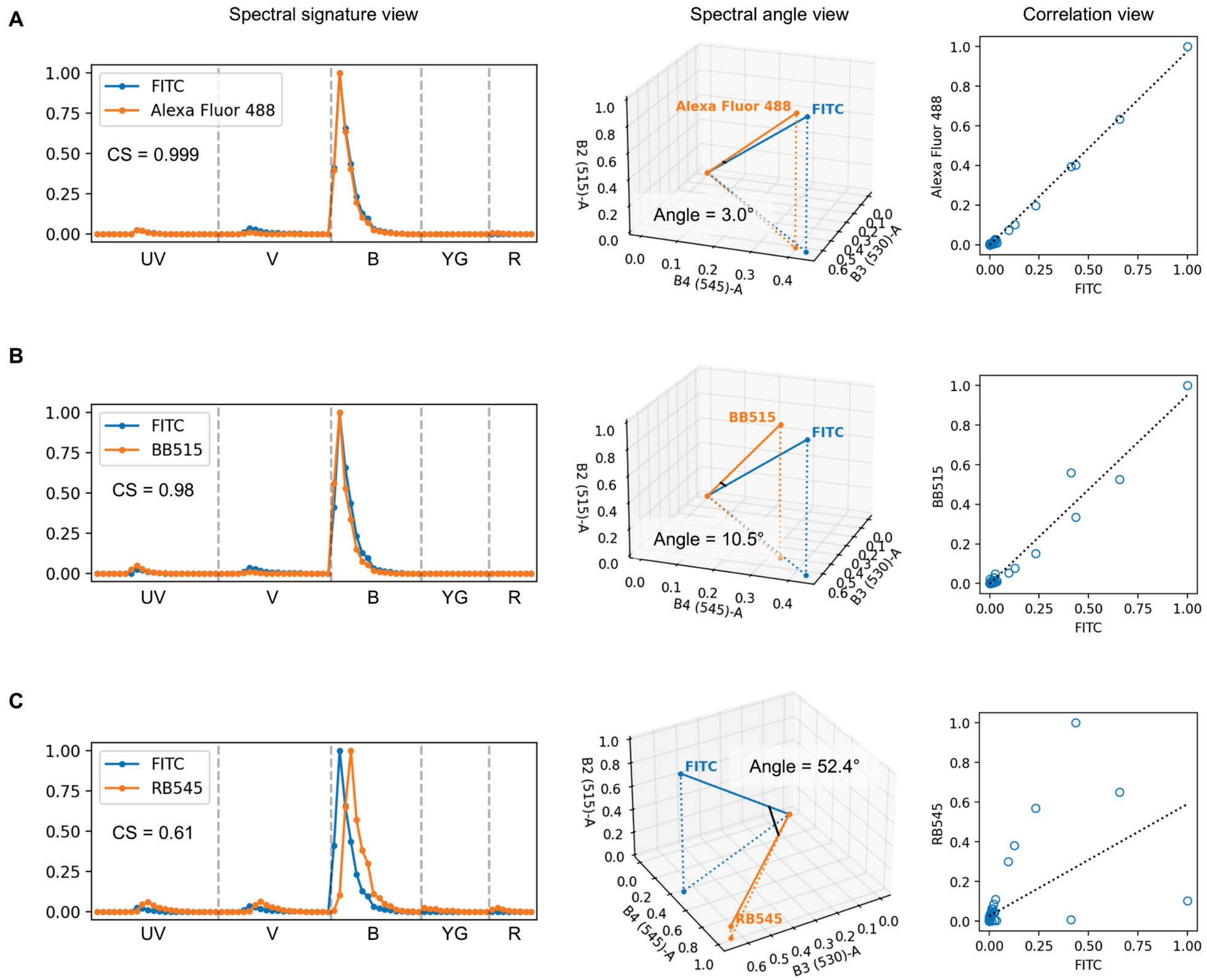

**SI Figure 1:** Visual explanation of cosine similarity (CS) in terms of overlapping normalized spectral signatures (left, "Spectral signature view"), angles between spectral vectors in high-dimensional space (middle, "Spectral angle view"), or correlation between spectral signatures (right, "Correlation view"). Example spectra, CS values, and spectral angles are shown on the BD FACSDiscover™ S8 for **(A)** a nearly identical fluorochrome pair, **(B)** a highly overlapping pair, and **(C)** a pair with moderate overlap. For the spectral angle view, vectors are shown in only three dimensions for illustration purposes, but angles and CS are calculated using the full 78-dimensional vectors. For the correlation view, each point corresponds to a single detector.

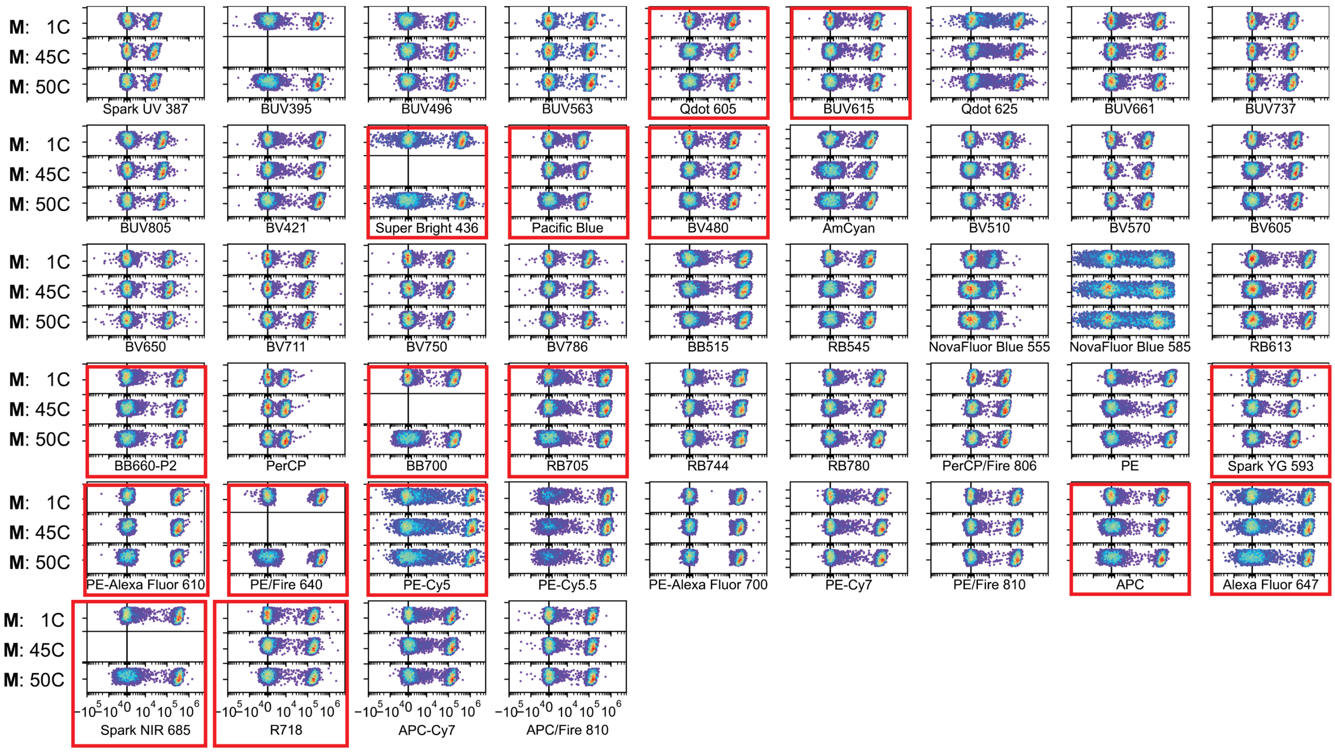

**SI Figure 2:** Visualization of USE for all fluorochromes (except viability and AF) in the 45C and 50C versions of OMIP-102. For each fluorochrome, a CD4 single-stain sample is unmixed using a single-color matrix containing that fluorochrome and AF (top subplot); the 45C matrix (middle subplot); and the full 50C matrix (bottom subplot). Vertical axis shows side scatter. Fluorochromes with USE exceeding 2 are outlined in red as in Figure 2C.

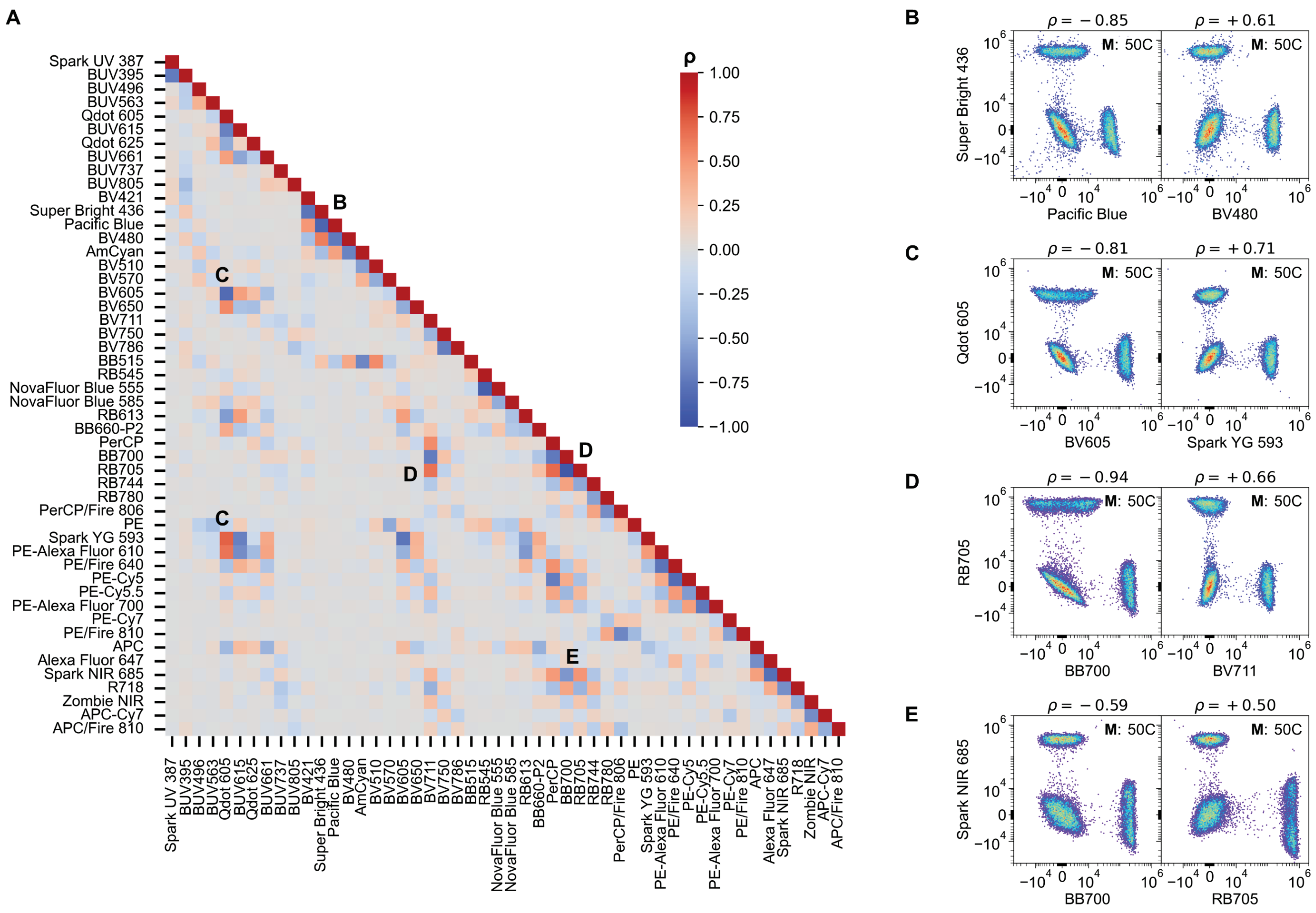

**SI Figure 3:** Correlation matrix and UDS Hallmark 2. **(A)** The presence of "tilted double-negative populations" in unmixed data can be quantified by calculating the correlation matrix for all unmixed parameters for an unstained population, shown here for OMIP-102. The correlation matrix is calculated by first computing the covariance matrix across all unmixed events, and subsequently dividing each entry of the covariance matrix by the product of the standard deviations (square root of diagonal covariance matrix entries) corresponding to that entry's row and column fluorochromes. Events with raw intensity above the 99.9th percentile or below the 0.01st percentile in any raw channel were excluded to remove the effect of outliers. **(B-E)** Examples of unmixed fluorochrome pairs from (A) with varying degrees of UDS in OMIP-102, manifesting as tilted double-negative populations. The visual severity and direction of tilt correspond to the magnitude and sign of the correlation coefficient ρ.

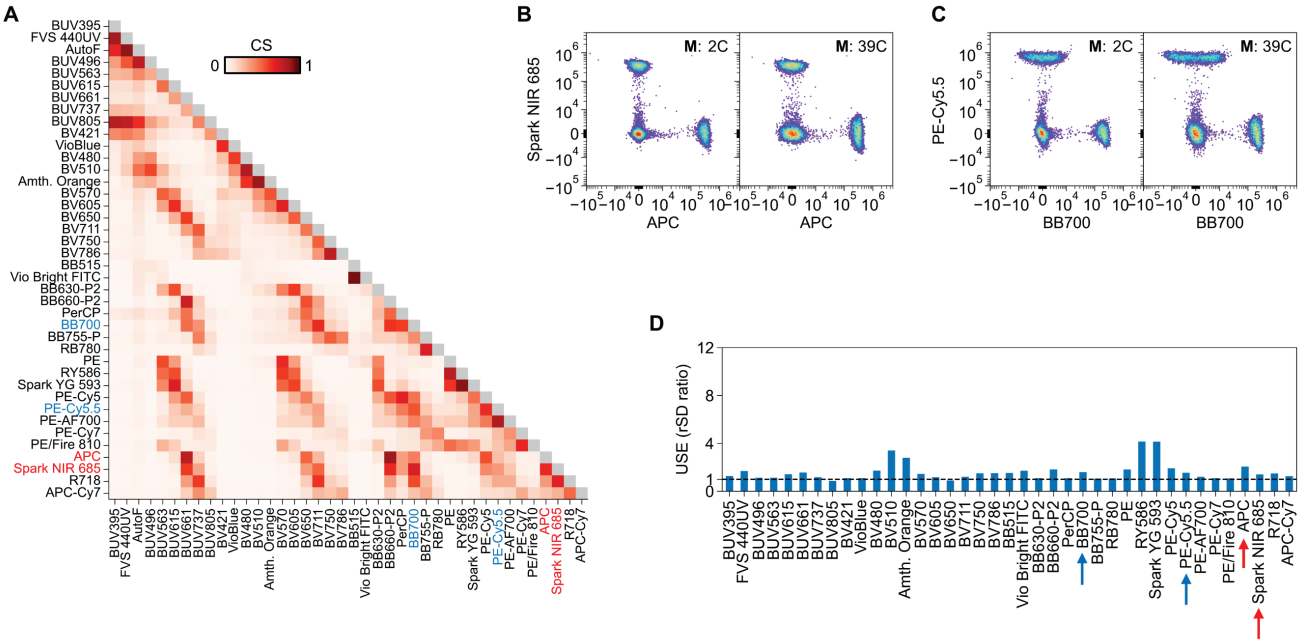

**SI Figure 4:** 39-color version of Panels 40C-A and 40C-B avoiding UDS in both the "APC" and "BB700" hotspots. **(A)** The matrix of cosine similarities shows identical values as Panels 40C-A and 40C-B for the common subset of fluorochromes included in the 39C version. **(B-C)** Minimal UDS is observed in the 39C panel unmixing (right) compared to 2C unmixing (left) for fluors in the "APC" (B) and "BB700" (C) hotspots. **(D)** Quantification of USE shows that fluorochromes in the "APC" and "BB700" spectral neighborhoods (arrows) are relatively unaffected by UDS compared to Panels 40C-A and 40C-B.

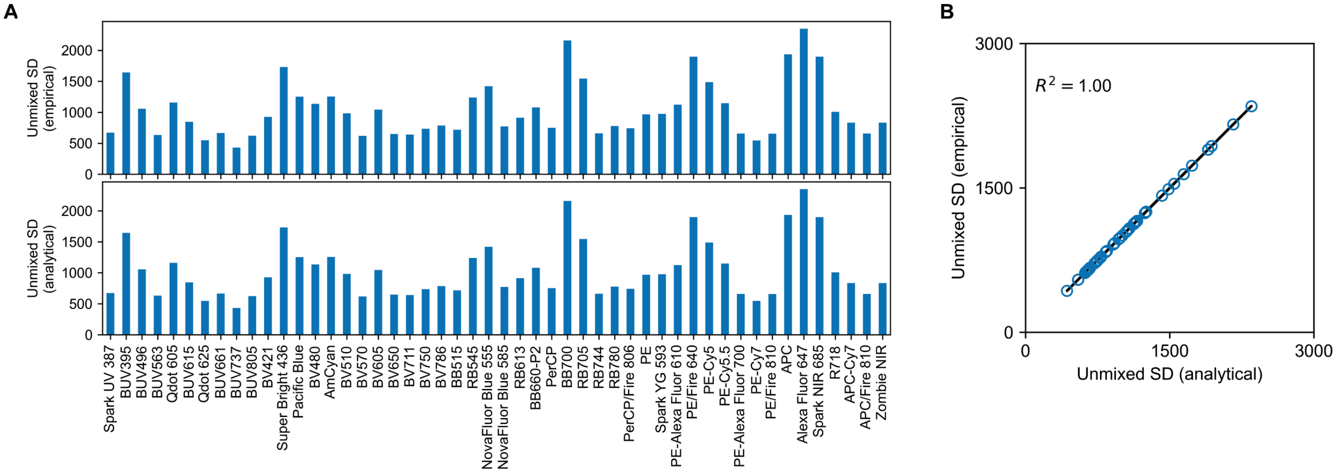

**SI Figure 5:** Agreement between calculations of spread from unmixed data or from raw covariance. **(A)** The unmixed standard deviation for each fluorochrome in OMIP-102 in an unstained sample is shown calculated empirically from unmixed data (top) or predicted analytically based on the raw covariance matrix and this panel's spectral matrix using Equation 3 (bottom). **(B)** Exact agreement (within floating-point rounding error) is observed between empirically calculated unmixed spread (y-axis) and unmixed spread predicted analytically from raw covariance (x-axis). For this example, calculations were performed on raw data with removed outliers (cells with intensities below the 1st percentile or above the 99th percentile in any raw fluorescence channel).

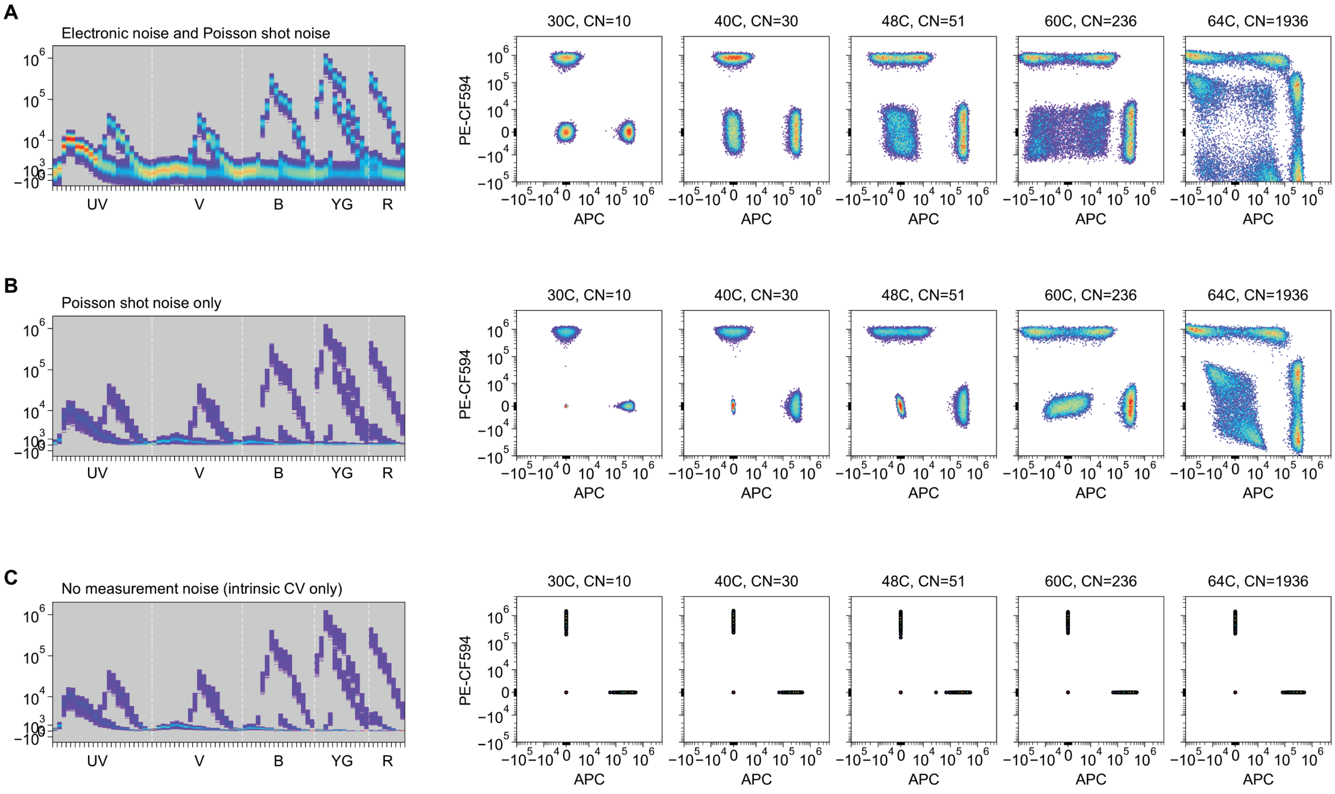

**SI Figure 6:** Simulated data with varying levels of raw measurement noise. Raw single-stain data (left) for the fluorochromes and panels described in Figure 1C was simulated using a linear mixture model based on measured fluorochrome spectral signatures and various sources of raw measurement noise. Each simulation includes three populations with 10,000 random events each: an unstained population (with AF), a PE-CF594+ population, and an APC+ population. Intensity of underlying AF and fluorochrome signals was modeled using gaussian distributions with a 20% intrinsic CV. Electronic noise was modeled by adding random values drawn from a zero-mean gaussian distribution with fixed variance across all detectors, while Poisson noise was modeled for each detector independently by sampling from a Poisson distribution with variance linearly proportional to the underlying signal for a given event. Simulated raw data was unmixed using OLS for the spectral matrices corresponding to the panels in Figure 1C (right). Noise sources included **(A)** detector electronic noise and Poisson shot noise, **(B)** Poisson shot noise alone, or **(C)** no added measurement noise. The magnitude and impact of UDS for a given panel is observed to vary depending on the magnitude of the raw measurement noise in (A) and (B). In the absence of any measurement noise (C), UDS is not observed, confirming that UDS only amplifies existing raw measurement noise.

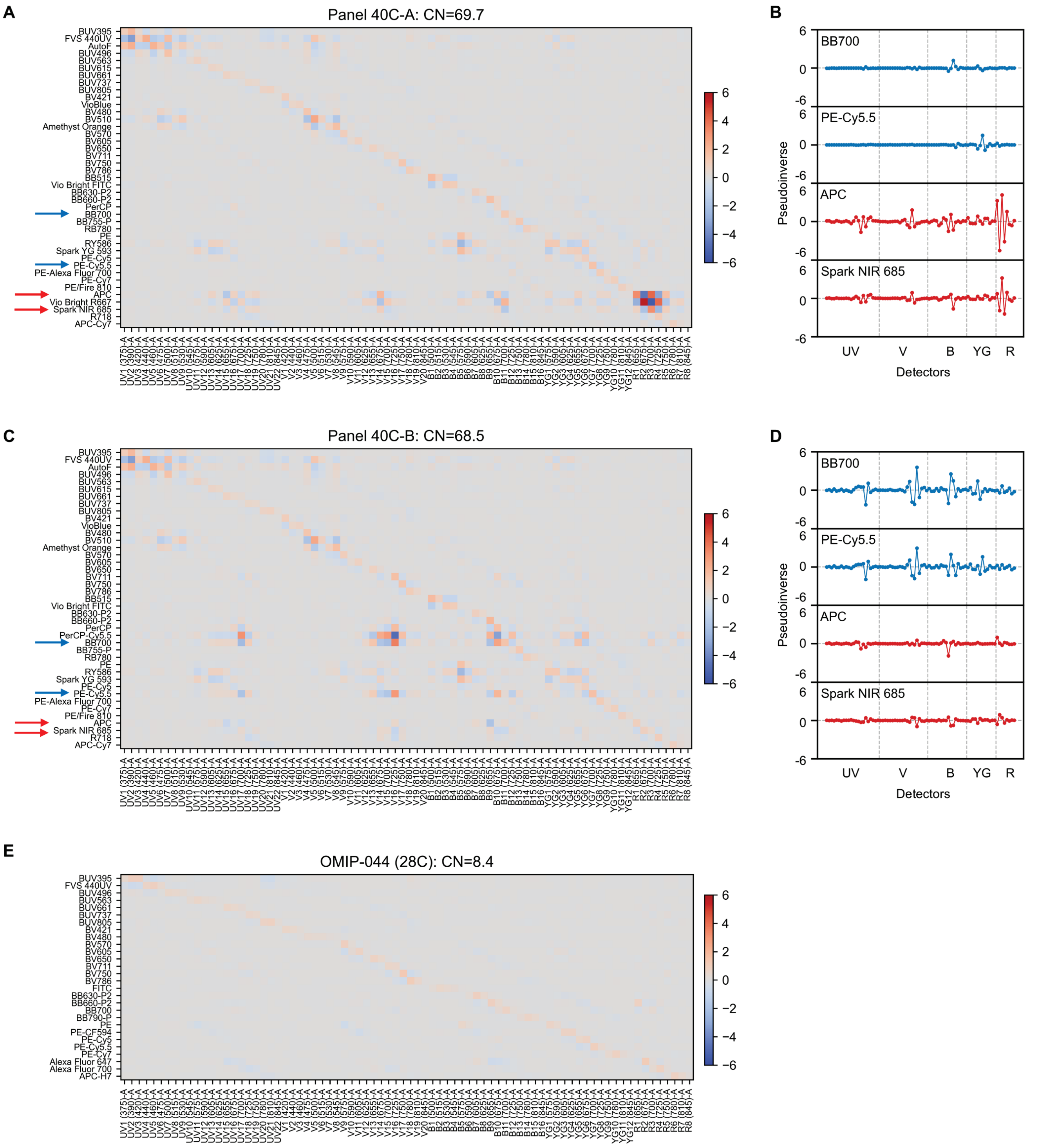

**SI Figure 7:** Spectral matrix pseudoinverse values vary for the same fluorochrome depending on the full panel. **(A)** The sign and magnitude of entries in the pseudoinverse matrix ("inverse spectra") are displayed on a heatmap for Panel 40C-A. Large-magnitude values with alternating positive and negative signs are observed for fluorochromes in the "APC" hotspot, especially around detectors on the red laser. **(B)** Line plots showing the inverse spectra of a subset of fluorochromes common to both panels. Large values are seen for fluorochromes in the "APC" hotspot when unmixed with Panel 40C-A. **(C)** Heatmap visualization of the inverse spectra for Panel 40C-B, showing large-magnitude values in the "BB700" hotspot. **(D)** Line plots showing the inverse spectra of the same fluorochromes as (B) when unmixed with Panel 40C-B. Large-magnitude values are observed in the "BB700" hotspot. **(E)** A heatmap of the inverse spectra for a well-conditioned panel with low spectral overlap, OMIP-044, shows no large oscillating values.

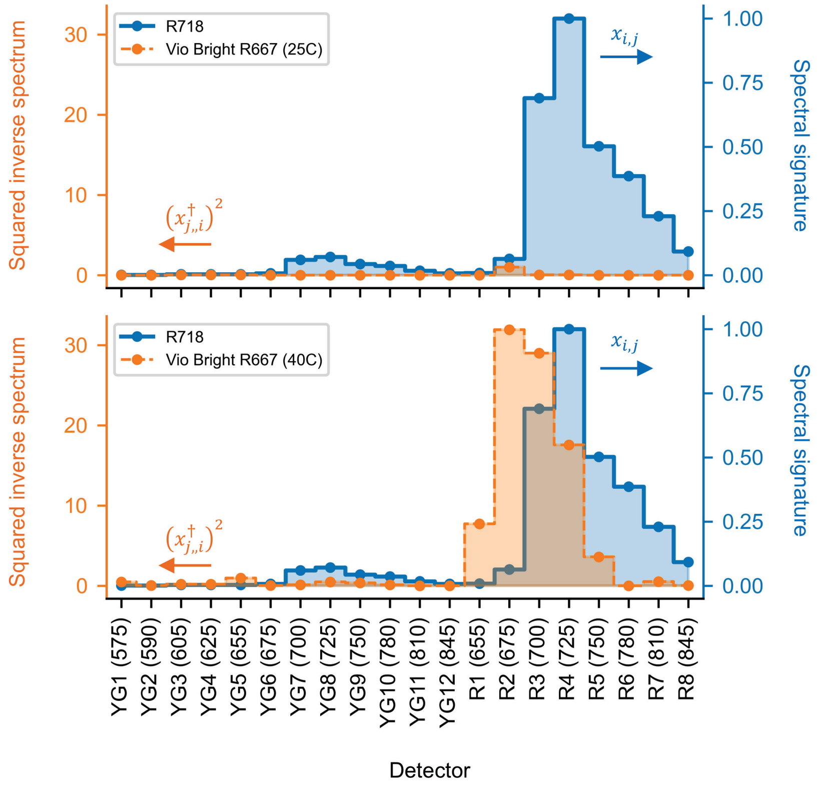

**SI Figure 8:** Explanation of UDS leading to increased SSE (Hallmark 3). The emission spectrum of R718 (blue, right axis) is displayed in comparison to the squared inverse spectrum of Vio Bright R667 (left axis) in the 25C backbone panel (top) or expanded 40C panel (bottom) corresponding to Figure 1B. Areas of overlap between the R718 spectrum and Vio Bright R667 inverse spectrum will lead to SSE from R718 into Vio Bright R667, as photonic shot noise from R718 emission is amplified proportionally to the squared inverse spectrum values of Vio Bright R667 (Equation SI-3). Due to collinearity in the 40C panel, the magnitude of entries in the 40C inverse spectrum of Vio Bright R667 corresponding to detectors with substantial R718 emission (e.g., R3-R4) are many orders of magnitude larger than the same entries in Vio Bright R667's 25C inverse spectrum. Data are shown for the BD FACSDiscover™ S8. Inverse spectra were calculated using all detectors, but only the subset corresponding to YG and R detectors, where the bulk of R718 emission occurs, are shown for clarity.

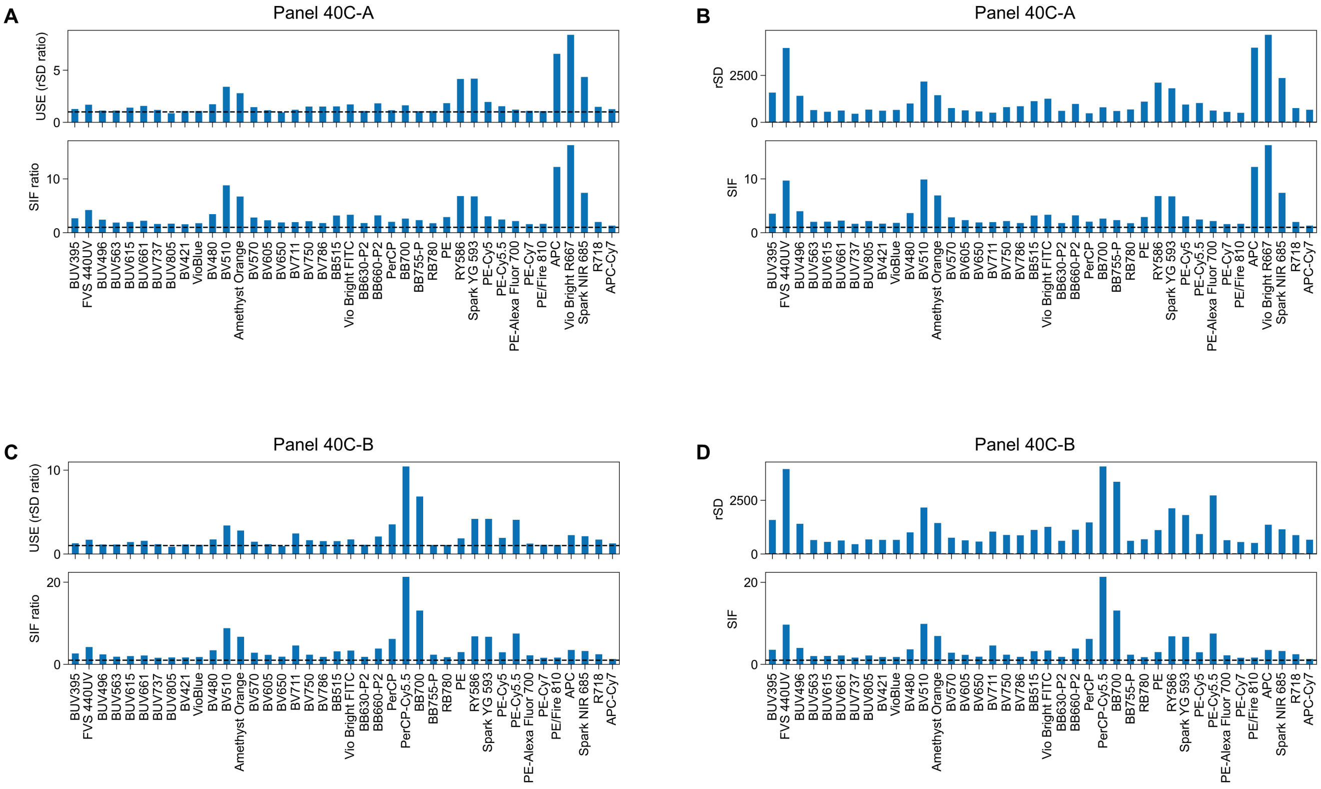

**SI Figure 9:** Comparison of Hotspot-derived metrics (SIFs and SIF ratios) to measured USE on a per-fluorochrome basis for Panels 40C-A and 40C-B. **(A)** USE as measured by the ratio of full-panel unmixed rSD to single-color unmixed rSD for an unstained population (top) is predicted closely by the ratio of full-panel SIFs to single-color SIFs for Panel 40C-A. The top and bottom values are plotted against each other in the inset of Figure 4C. **(B)** Similar relationships can be observed between the rSD of the full-panel unmixed sample (same values as in (A), but not divided by single-color unmixed rSD) and the SIFs. Different patterns of spread between the ratiometric values (A) and non-ratiometric values (B) are primarily due to AF, which affects both the measured and predicted UDS for a given fluorochrome depending on that fluorochrome's spectral similarity to AF, even in the context of a single-color matrix containing only that fluorochrome and AF. **(C)** and **(D)** show the same analysis as (A) and (B) but for Panel 40C-B.

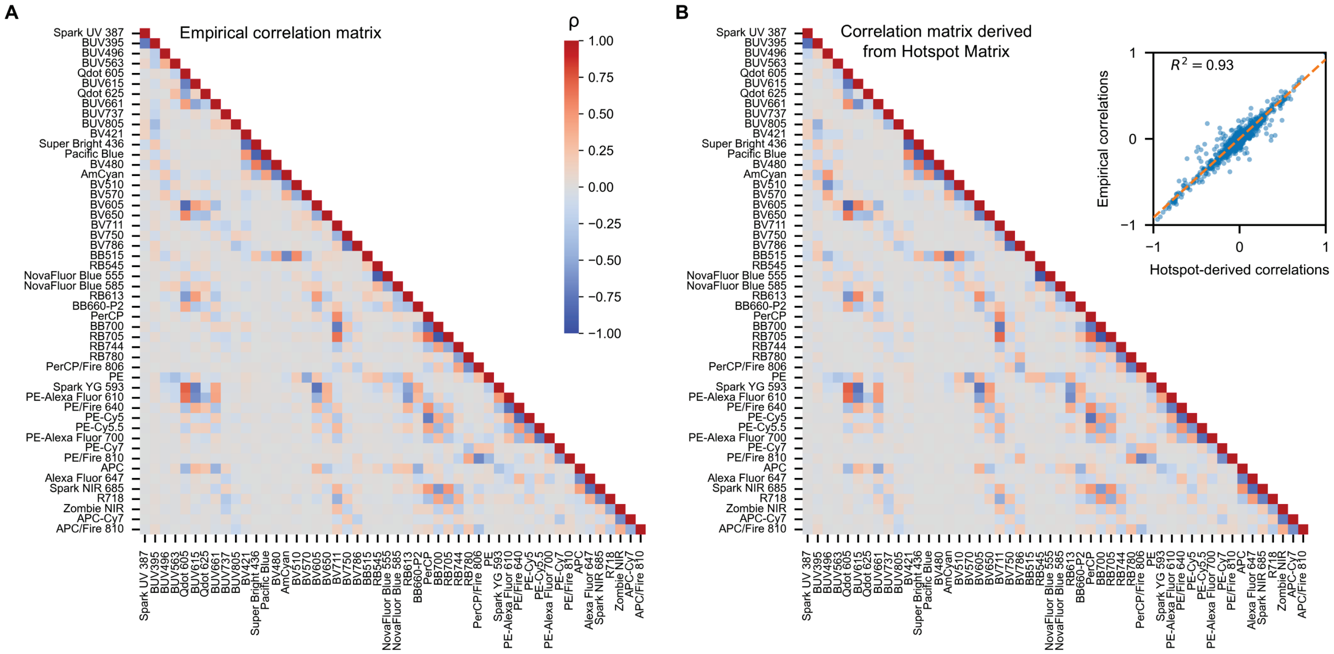

**SI Figure 10:** Predicting measured correlation (Hallmark 2) with a modified Hotspot Matrix. **(A)** Empirical correlation matrix for unmixed parameters in OMIP-102 as calculated in SI Figure 3. Correlation coefficients describe the direction and severity of tilt in double-negative populations. **(B)** Because the Hotspot Matrix's precursor matrix $(X^T X)^{-1}$ is nominally proportional to the unmixed covariance matrix, it can be used to calculate a predicted correlation matrix through the same scaling process described in SI Figure 3. The resulting predicted correlation matrix closely matches the empirical correlation matrix in (A) (inset).

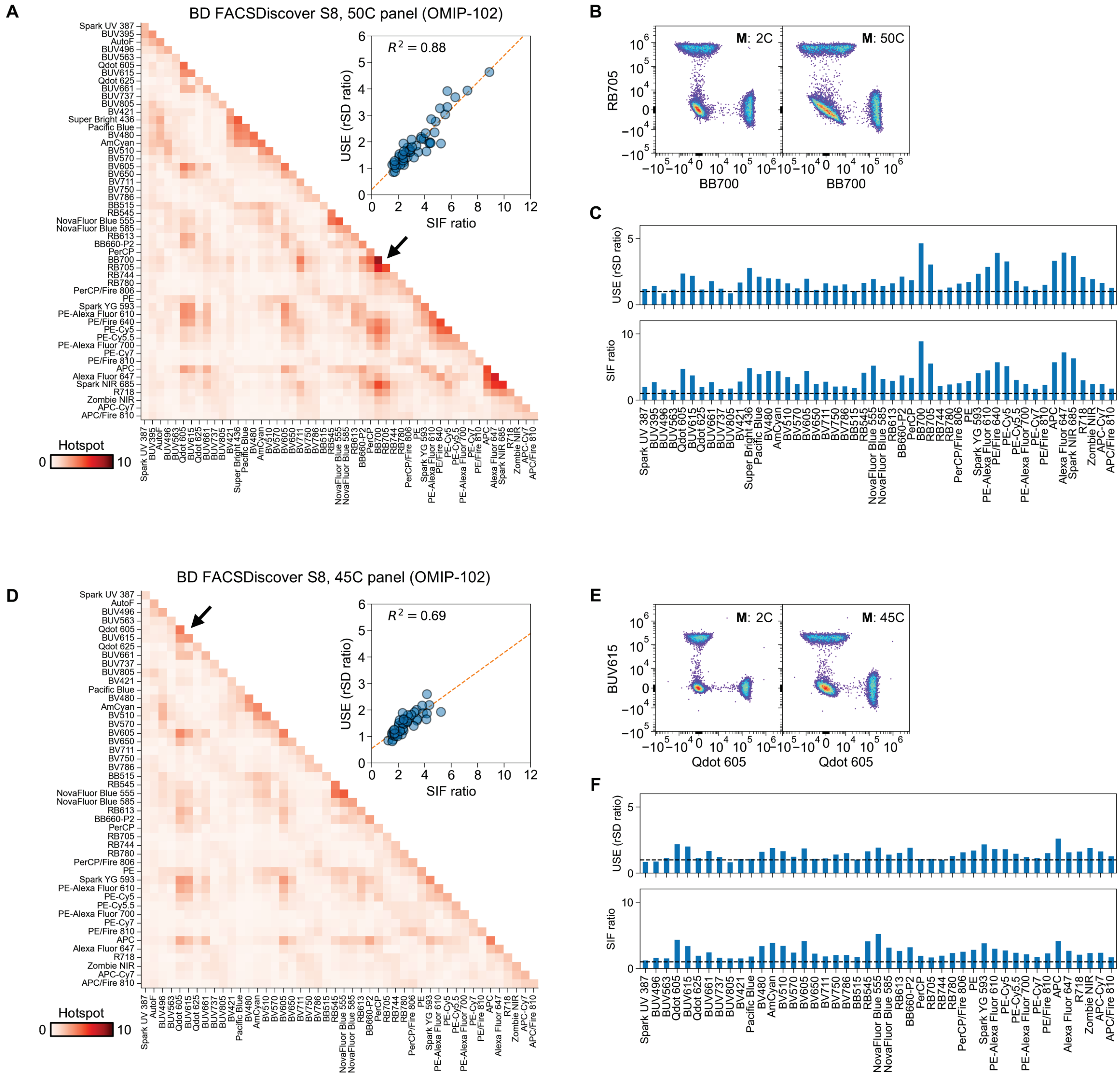

**SI Figure 11:** Hotspot Matrix and USE for OMIP-102 on the BD FACSDiscover™ S8. **(A)** Hotspot Matrix and comparison of USE and SIF ratios for the 50C version of OMIP-102, as measured on the BD FACSDiscover™ S8. **(B)** Example unmixing for a pair of fluorochromes in a hotspot, indicated by the arrow in (A). **(C)** Comparison of USE and SIF ratios on a per-fluorochrome basis. **(D-F)** Similar analysis as A-C for the 45C version of OMIP-102.

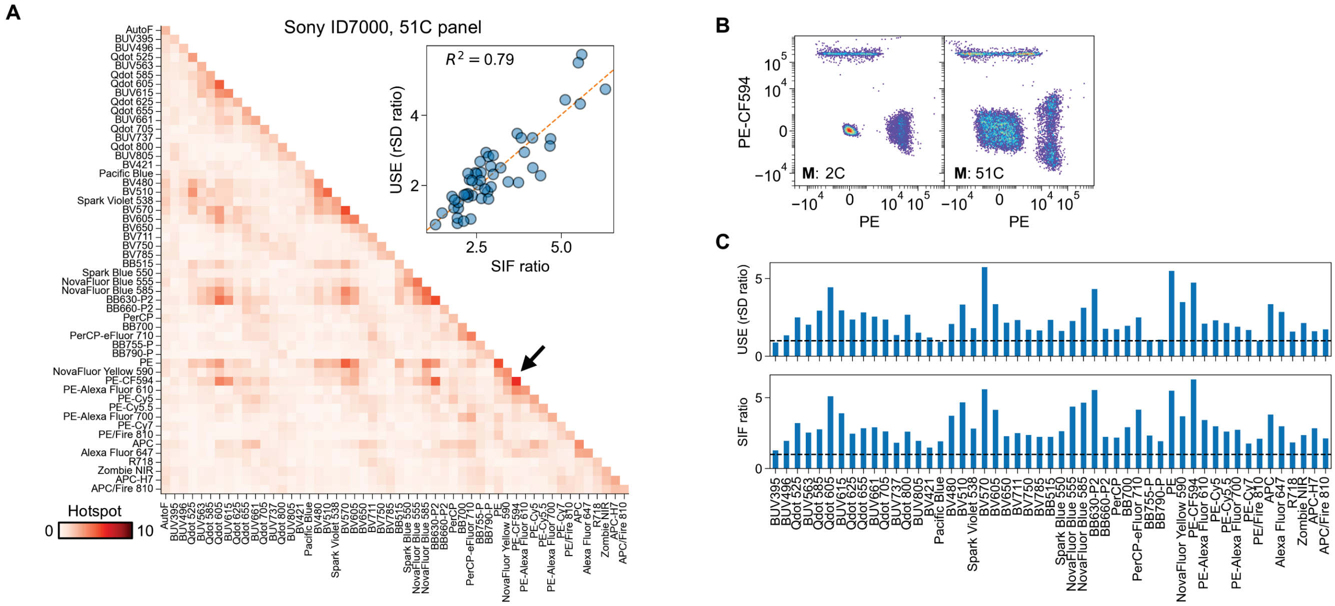

**SI Figure 12:** Hotspot Matrix and USE for a 51C panel on the Sony ID7000™. **(A)** Hotspot Matrix and comparison of USE and SIF ratios for the 51C panel shown in Figure 1A, as measured on the Sony ID7000™. **(B)** Example unmixing for a pair of fluorochromes in a hotspot, indicated by the arrow in (A). For this panel, single-stain bead controls are shown. **(C)** Comparison of USE and SIF ratios on a per-fluorochrome basis.

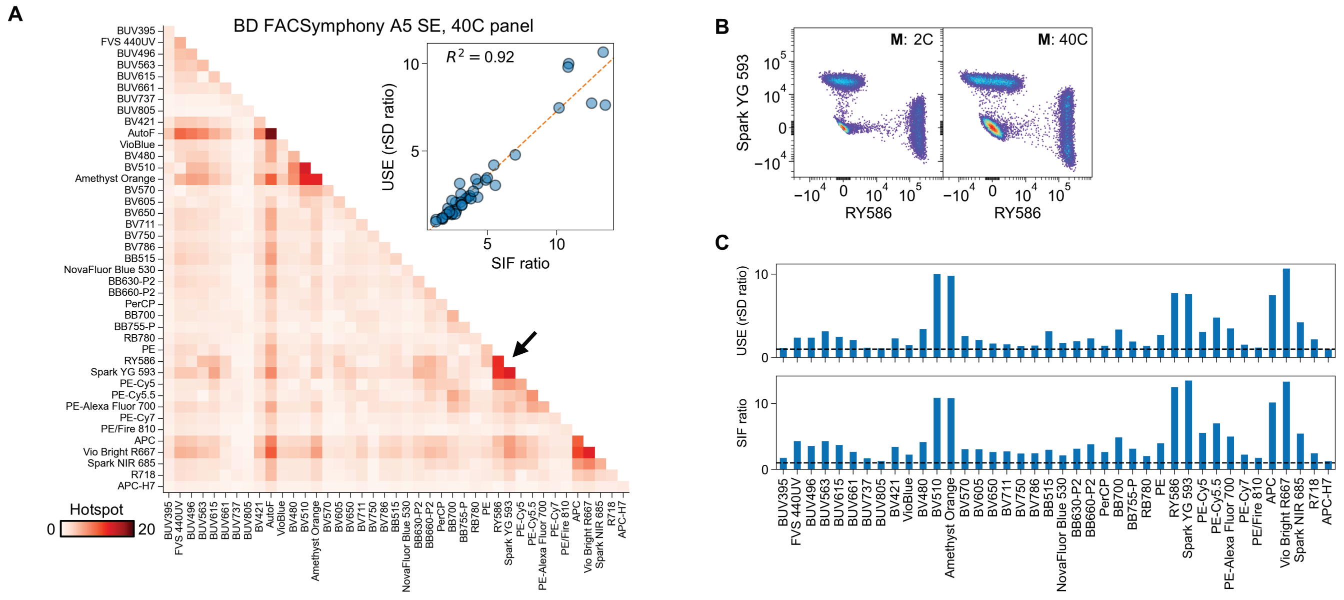

**SI Figure 13:** Hotspot Matrix and USE for a 40C panel on the BD FACSymphony™ A5 SE. **(A)** Hotspot Matrix and comparison of USE and SIF ratios for the 40C panel shown in Figure 1, as measured on the BD FACSymphony™ A5 SE. **(B)** Example unmixing for a pair of fluorochromes in a hotspot, indicated by the arrow in (A). **(C)** Comparison of USE and SIF ratios on a per-fluorochrome basis.

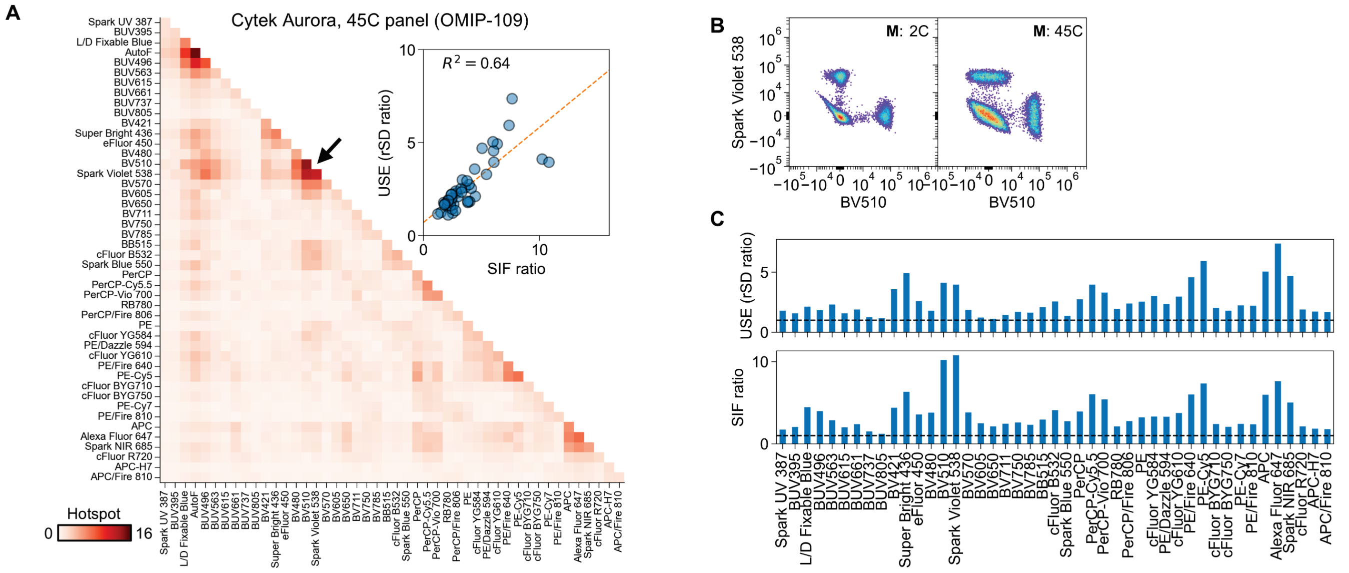

**SI Figure 14:** Hotspot Matrix and USE for OMIP-109 on the Cytek® Aurora[2]. **(A)** Hotspot Matrix and comparison of USE and SIF ratios for OMIP-109 on the Cytek® Aurora. **(B)** Example unmixing for a pair of fluorochromes in a hotspot, indicated by the arrow in (A). **(C)** Comparison of USE and SIF ratios on a per-fluorochrome basis.

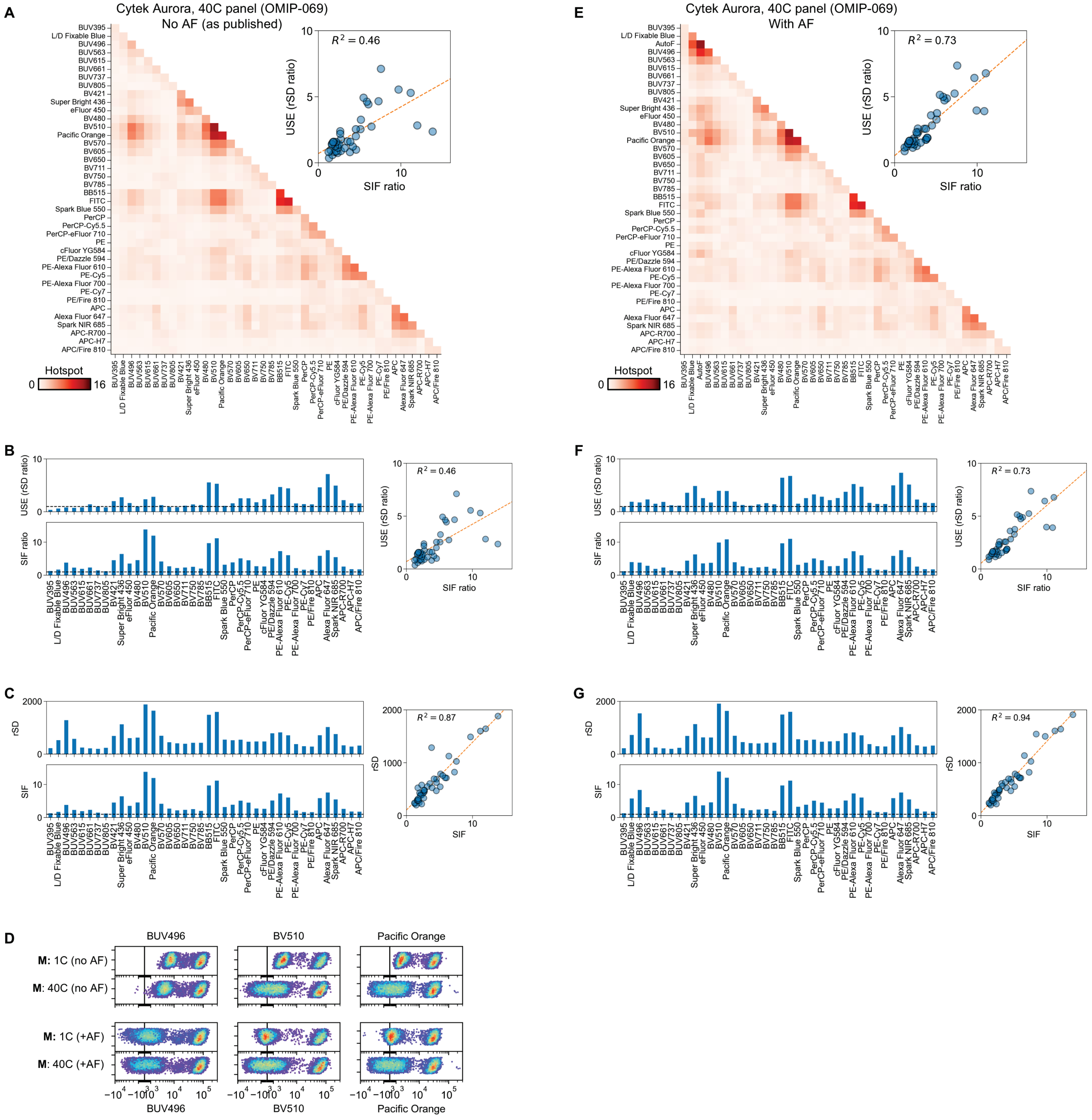

**SI Figure 15:** Effect of AF unmixing on Hotspot analysis and USE. **(A)** Hotspot Matrix for OMIP-069 on the Cytek® Aurora, with AF excluded from unmixing as originally published[3]. While problematic areas of UDS are still clearly identified, SIF ratios have some discrepancies from measured USE (inset). **(B)** Examination of USE (top) and SIF ratios (bottom) by fluorochrome reveals that discrepancies are observed for fluorochromes with substantial spectral overlap with AF, such as BUV496, BV510, and Pacific Orange. Some fluorochromes have USE values below 1, indicating that UDS is higher for those fluorochromes when unmixed using a single-color matrix as opposed to the 40C matrix, when AF is excluded from both. This likely occurs because intrinsic AF intensity variation across cells contributes to different fluorochromes' unmixed variances in an uncontrolled manner when

AF is excluded from unmixing. **(C)** Non-ratio-based metrics that exclude the behavior of single-color unmixing show a stronger agreement (right), although outliers are still present. **(D)** Comparison of single-color and full-panel unmixing without AF (top) reveals that the unaccounted-for signal from AF results in artificial elevation of unmixed MFI for a subset of fluorochromes, an indication that the real AF signal cannot be correctly modeled by a spectral matrix without AF. By contrast, inclusion of AF (bottom) shows expected extraction of AF signal from each unmixed fluorochrome channel. **(E-G)** Hotspot Matrix and associated metrics for OMIP-069 with inclusion of AF. Although the correspondence between USE and SIF ratios is improved (E, inset), the inclusion of AF itself introduces additional UDS in some fluorochromes compared to the AF-free matrix.

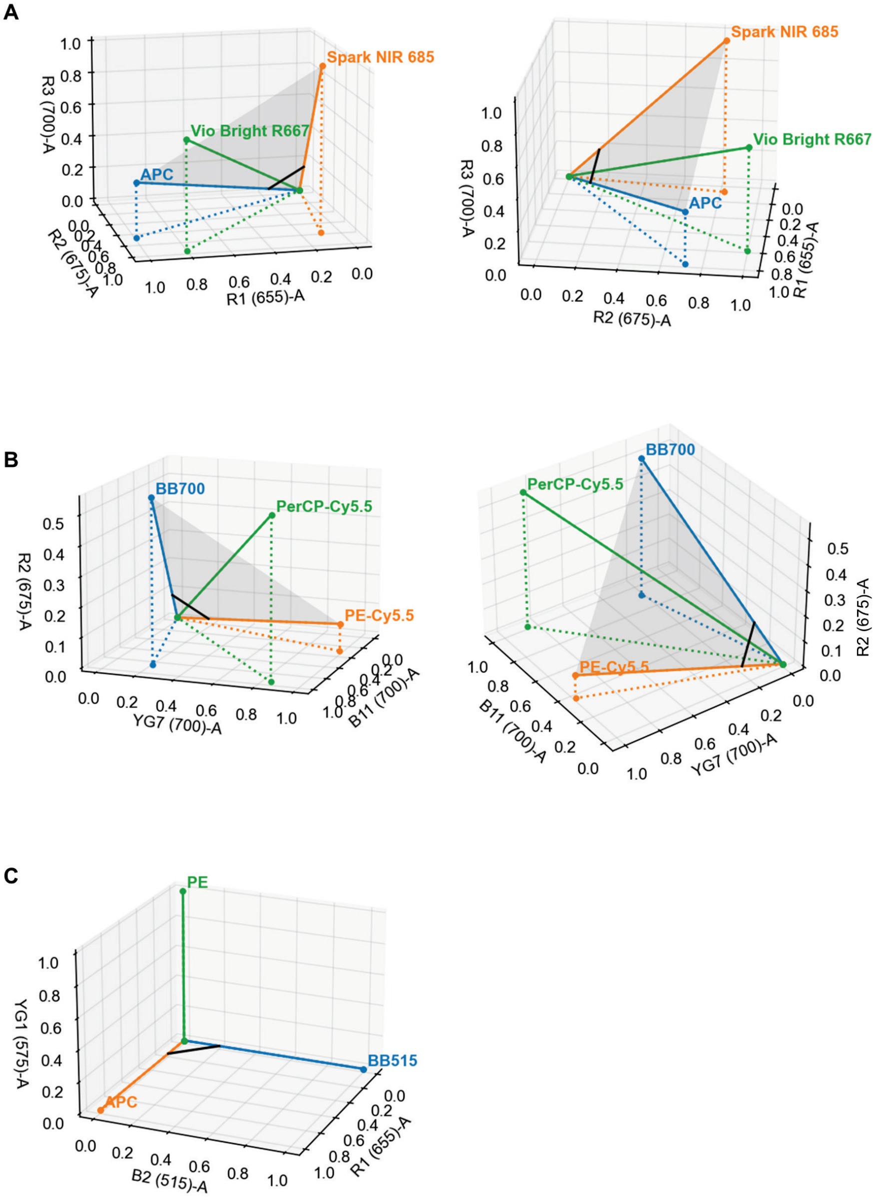

**SI Figure 16:** Visual illustration of collinear spectra. Spectral signatures in the "APC" and "BB700" hotspots from Panels 40C-A and 40C-B are plotted as three-dimensional vectors in **(A)** and **(B)**, where the plot axes correspond to three detectors near the emission peaks for each fluorochrome. Although each pair of fluorochromes within the hotspot is separated by a modest spectral angle, all three vectors fall nearly in the same two-dimensional plane (gray), indicating collinearity. Multiple views of each plot are shown from varying angles to illustrate the vectors' positions relative to each other in three dimensions. A highly non-collinear fluorochrome combination is shown in **(C)** for comparison.

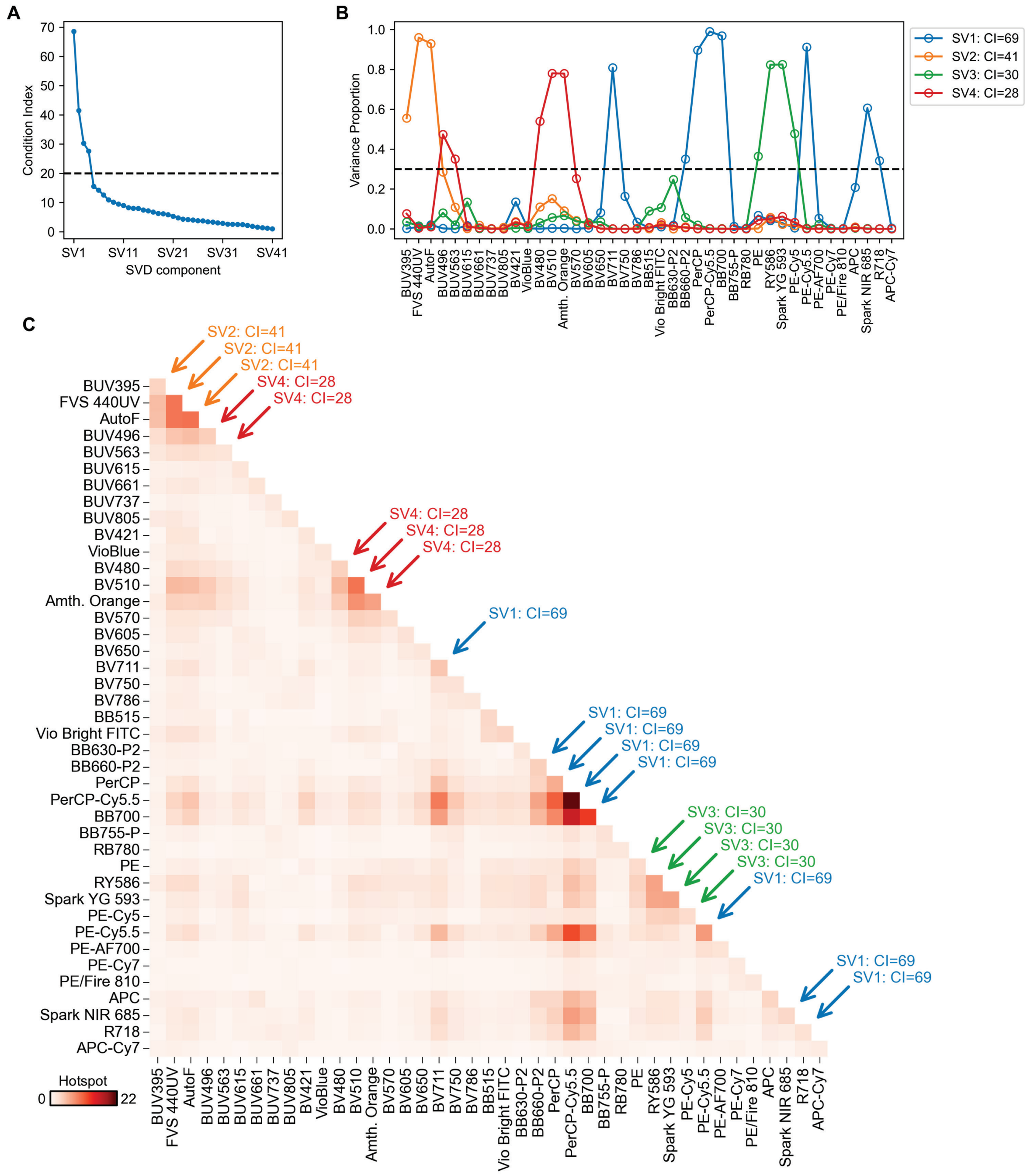

**SI Figure 17:** Example of Variance Decomposition Proportion (VDP) analysis for Panel 40C-B. **(A)** Singular value decomposition (SVD) is performed on the panel's spectral matrix, and the condition index (CI) of each component of the SVD (SV1, SV2, SV3, etc.) is computed by taking the ratio of the largest singular value to that component's singular value. Components of the SVD with a CI above an empirically selected threshold (here, CI=20) indicate the presence of collinearity. **(B)** The proportion of each fluorochrome's predicted unmixed variance that is attributable to each component of the SVD is computed following the method of Belsley[4–6]. Any set of fluorochromes that have a high proportion of their variance attributable to the same high-CI SVD component are considered collinear. Here, a variance proportion threshold of 0.3 (30% of variance) is chosen as an empirical lower limit indicating collinearity. **(C)** The Hotspot Matrix for Panel 40C-B is overlaid with labels indicating the high-CI

SVD components with which each fluorochrome is associated, if any. Fluorochromes associated with the same high-CI SVD components have some degree of collinearity with each other. VDP analysis provides unambiguous confirmation that PE-Cy5.5 and BV711 are part of the same hotspot as BB700 and PerCP-Cy5.5 in this panel, since all are associated with SV1. It also reveals that BUV496 is more strongly associated with the BV510 / Amethyst Orange hotspot (SV4), while also being more weakly associated with the hotspot around AF and Fixable Viability Stain 440UV (SV2). We note that the choice of CI and variance proportion thresholds will affect which specific collinear combinations are highlighted via VDP, and selection of appropriate thresholds is a common challenge in VDP analysis[5].

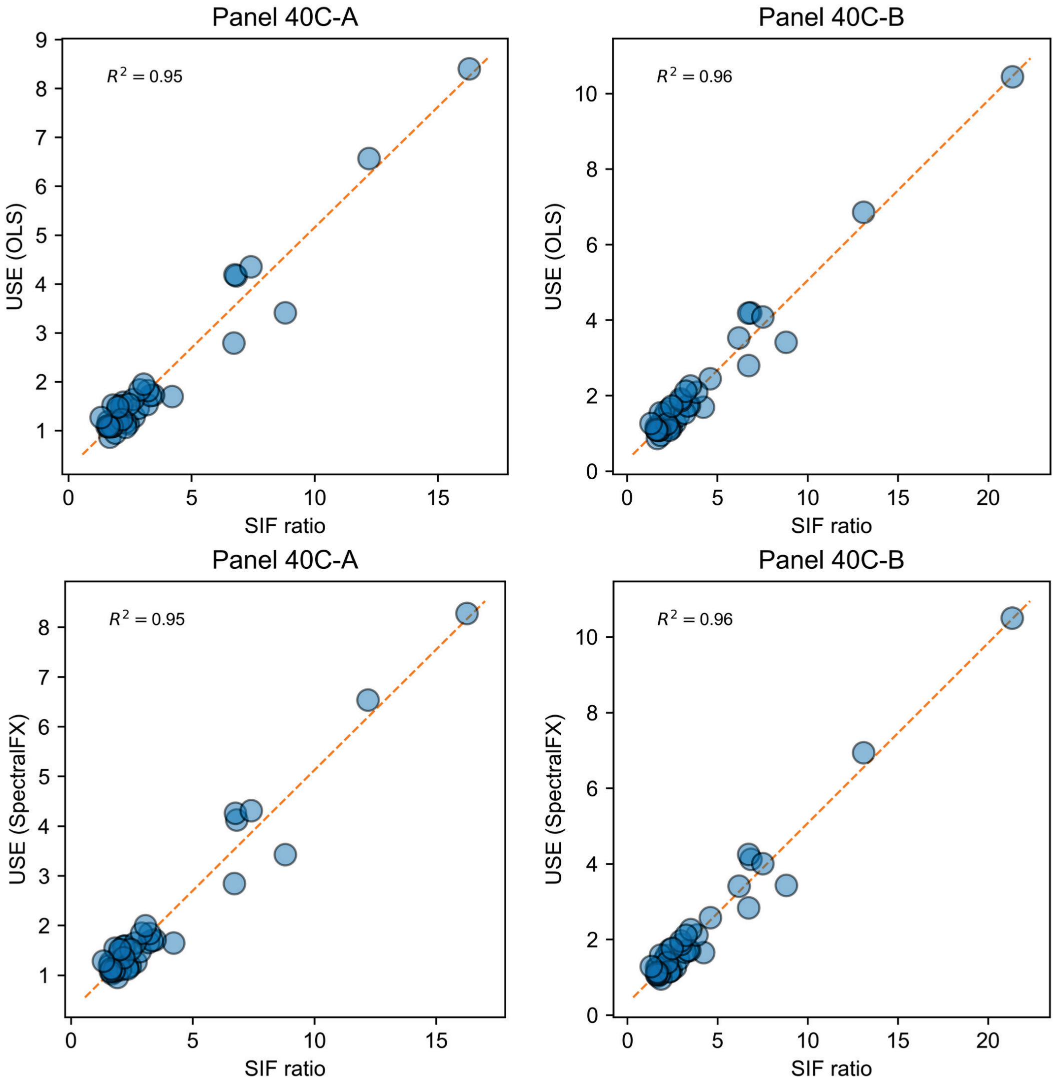

**SI Figure 18:** Using SIFs to predict USE for OLS and non-OLS unmixing methods. USE was calculated by taking the ratio of unmixed rSD for an unstained sample between full-panel unmixing and single-color unmixing for each unmixed parameter in panels 40C-A (left) and 40C-B (right). Unmixing was performed using either OLS unmixing (top) or BD SpectralFX™ unmixing (bottom), a proprietary non-OLS algorithm. SIFs derived from the Hotspot Matrix serve as a strong predictor of USE for both unmixing methods.

### SUPPLEMENTARY TABLES

| Figure | Fig. 1A | Fig. 1A | Fig. 1B, D-F | Fig. 1B, D-F | Fig. 2 | Fig. 2 | Figs. 3-4, 5A | Figs. 3-4 | Fig. 5B | Fig. 5C | Fig. 5D | Fig. 5E | Fig. 5F |
|---|---|---|---|---|---|---|---|---|---|---|---|---|---|
| Description | *Backbone* | *Full panel* | *Backbone* | *Full panel* | *OMIP-102 50C* | *OMIP-102 45C* | *Panel 40C-A* | *Panel 40C-B* | *Panel 40C-A2* | *Panel 40C-A3* | *Panel 40C-A4* | *Panel 40C-A5* | *Panel 45C-A6* |
| Instrument | *ID7000* | *ID7000* | *S8* | *S8* | *S8* | *S8* | *S8* | *S8* | *S8* | *S8* | *S8* | *S8* | *S8* |
| Panel Size | *20C+AF* | *51C+AF* | *25C+AF* | *40C+AF* | *50C+AF* | *45C+AF* | *40C+AF* | *40C+AF* | *40C+AF* | *40C+AF* | *40C+AF* | *40C+AF* | *45C+AF* |
| | | Qdot 525 | BUV395 | BUV395 | Spark UV 387 | Spark UV 387 | | | | | | | Spark UV 387 |
| | | Qdot 585 | FVS440UV | FVS440UV | BUV395 | | BUV395 | BUV395 | BUV395 | BUV395 | BUV395 | BUV395 | BUV395 |
| | | Qdot 605 | | BUV496 | BUV496 | BUV496 | FVS440UV | FVS440UV | FVS440UV | FVS440UV | FVS440UV | | |
| | | Qdot 625 | BUV563 | BUV563 | BUV563 | BUV563 | BUV496 | BUV496 | BUV496 | BUV496 | BUV496 | BUV496 | BUV496 |
| | | Qdot 655 | BUV615 | BUV615 | Qdot 605 | Qdot 605 | BUV563 | BUV563 | BUV563 | BUV563 | BUV563 | BUV563 | BUV563 |
| | | Qdot 705 | BUV661 | BUV661 | BUV615 | BUV615 | BUV615 | BUV615 | BUV615 | BUV615 | BUV615 | BUV615 | BUV615 |
| | Qdot 800 | Qdot 800 | BUV737 | BUV737 | Qdot 625 | Qdot 625 | | | | Qdot 625 | Qdot 625 | Qdot 625 | Qdot 625 |
| | | BUV395 | BUV805 | BUV805 | BUV661 | BUV661 | BUV661 | BUV661 | BUV661 | BUV661 | BUV661 | BUV661 | BUV661 |
| | | BUV496 | BV421 | BV421 | BUV737 | BUV737 | BUV737 | BUV737 | BUV737 | BUV737 | BUV737 | BUV737 | BUV737 |
| | BUV563 | BUV563 | VioBlue | VioBlue | BUV805 | BUV805 | BUV805 | BUV805 | BUV805 | BUV805 | BUV805 | BUV805 | BUV805 |
| | BUV615 | BUV615 | | BV480 | BV421 | BV421 | BV421 | BV421 | BV421 | BV421 | BV421 | BV421 | BV421 |
| | | BUV661 | BV510 | BV510 | SuBr 436 | | VioBlue | VioBlue | VioBlue | VioBlue | VioBlue | VioBlue | VioBlue |
| | | BUV737 | | Amth. Orange | Pacific Blue | Pacific Blue | BV480 | BV480 | BV480 | BV480 | BV480 | BV480 | BV480 |
| | BUV805 | BUV805 | BV570 | BV570 | BV480 | BV480 | BV510 | BV510 | BV510 | BV510 | BV510 | BV510 | BV510 |
| | | BV421 | | BV605 | AmCyan | AmCyan | Amth. Orange | Amth. Orange | Amth. Orange | | | | |
| | Pacific Blue | Pacific Blue | | BV650 | BV510 | BV510 | BV570 | BV570 | BV570 | BV570 | BV570 | BV570 | BV570 |
| | | BV480 | | BV711 | BV570 | BV570 | BV605 | BV605 | BV605 | BV605 | BV605 | BV605 | BV605 |
| | BV510 | BV510 | | BV750 | BV605 | BV605 | BV650 | BV650 | BV650 | BV650 | BV650 | BV650 | BV650 |
| | Spark Violet 538 | Spark Violet 538 | | BV786 | BV650 | BV650 | BV711 | BV711 | BV711 | BV711 | BV711 | BV711 | BV711 |
| | BV570 | BV570 | BB515 | BB515 | BV711 | BV711 | BV750 | BV750 | BV750 | BV750 | BV750 | BV750 | BV750 |
| | BV605 | BV605 | NFB 530 | NFB 530 | BV750 | BV750 | BV786 | BV786 | BV786 | BV786 | BV786 | BV786 | BV786 |
| | | BV650 | | BB630-P2 | BV786 | BV786 | BB515 | BB515 | BB515 | BB515 | BB515 | BB515 | BB515 |
| | | BV711 | BB660-P2 | BB660-P2 | BB515 | BB515 | Vio Bright FITC | Vio Bright FITC | Vio Bright FITC | Vio Bright FITC | Vio Bright FITC | Vio Bright FITC | Vio Bright FITC |
| | | BV750 | PerCP | PerCP | RB545 | RB545 | | | | | | | NFB 585 |
| | | BV786 | | BB700 | NFB 555 | NFB 555 | BB630-P2 | BB630-P2 | BB630-P2 | BB630-P2 | BB630-P2 | BB630-P2 | BB630-P2 |
| | | BB515 | | BB755-P | NFB 585 | NFB 585 | BB660-P2 | BB660-P2 | BB660-P2 | BB660-P2 | BB660-P2 | BB660-P2 | BB660-P2 |
| | Spark Blue 550 | Spark Blue 550 | RB780 | RB780 | RB613 | RB613 | PerCP | PerCP | PerCP | PerCP | PerCP | PerCP | PerCP |
| | | NFB 555 | | PE | BB660-P2 | BB660-P2 | | PerCP-Cy5.5 | | | | | |
| | NFB 585 | NFB 585 | RY586 | RY586 | PerCP | PerCP | BB700 | BB700 | BB700 | BB700 | BB700 | BB700 | BB700 |
| | | BB630-P2 | Spark YG 593 | Spark YG 593 | BB700 | | BB755-P | BB755-P | BB755-P | BB755-P | BB755-P | BB755-P | BB755-P |
| | PerCP | PerCP | PE-Cy5 | PE-Cy5 | RB705 | RB705 | RB780 | RB780 | RB780 | RB780 | RB780 | RB780 | RB780 |
| | | BB660-P2 | PE-Cy5.5 | PE-Cy5.5 | RB744 | RB744 | | | | | | | PerCP/Fire 806 |
| | | BB700 | PE-AF700 | PE-AF700 | RB780 | RB780 | PE | PE | PE | PE | PE | PE | PE |
| | PerCP-eF710 | PerCP-eF710 | PE-Cy7 | PE-Cy7 | PerCP/Fire 806 | PerCP/Fire 806 | RY586 | RY586 | RY586 | RY586 | RY586 | RY586 | RY586 |
| | | BB755 | PE/Fire 810 | PE/Fire 810 | PE | PE | Spark YG 593 | Spark YG 593 | Spark YG 593 | Spark YG 593 | RY610 | RY610 | RY610 |
| | | BB790 | | APC | Spark YG 593 | Spark YG 593 | | | | | | | PE-AF610 |
| | | PE | | Spark NIR 685 | PE-AF610 | PE-AF610 | PE-Cy5 | PE-Cy5 | PE-Cy5 | PE-Cy5 | PE-Cy5 | PE-Cy5 | PE-Cy5 |
| | NFB 590 | NFB 590 | Vio Bright R667 | Vio Bright R667 | PE/Fire 640 | | PE-Cy5.5 | PE-Cy5.5 | PE-Cy5.5 | PE-Cy5.5 | PE-Cy5.5 | PE-Cy5.5 | PE-Cy5.5 |
| | | PE-CF594 | R718 | R718 | PE-Cy5 | PE-Cy5 | PE-AF700 | PE-AF700 | PE-AF700 | PE-AF700 | PE-AF700 | PE-AF700 | PE-AF700 |
| | | PE-AF610 | | APC-H7 | PE-Cy5.5 | PE-Cy5.5 | | | | | | | RY743 |
| | PE-Cy5 | PE-Cy5 | | | PE-AF700 | PE-AF700 | PE-Cy7 | PE-Cy7 | PE-Cy7 | PE-Cy7 | PE-Cy7 | PE-Cy7 | PE-Cy7 |
| | | PE-Cy5.5 | | | PE-Cy7 | PE-Cy7 | PE/Fire 810 | PE/Fire 810 | PE/Fire 810 | PE/Fire 810 | PE/Fire 810 | PE/Fire 810 | PE/Fire 810 |
| | PE-AF700 | PE-AF700 | | | PE/Fire 810 | PE/Fire 810 | APC | APC | APC | APC | APC | APC | APC |
| | | PE-Cy7 | | | APC | APC | Spark NIR 685 | Spark NIR 685 | Spark NIR 685 | Spark NIR 685 | Spark NIR 685 | Spark NIR 685 | Spark NIR 685 |
| | PE/Fire 810 | PE/Fire 810 | | | AF647 | AF647 | Vio Bright R667 | | | | | | |
| | | APC | | | Spark NIR 685 | | R718 | R718 | R718 | R718 | R718 | R718 | R718 |
| | AF647 | AF647 | | | R718 | R718 | | | | | | Zombie NIR | Zombie NIR |
| | | R718 | | | APC-Cy7 | APC-Cy7 | APC-Cy7 | APC-Cy7 | APC-Cy7 | APC-Cy7 | APC-Cy7 | APC-Cy7 | APC-Cy7 |
| | Zombie NIR | Zombie NIR | | | APC/Fire 810 | APC/Fire 810 | | | APC/Fire 810 | APC/Fire 810 | APC/Fire 810 | APC/Fire 810 | APC/Fire 810 |
| | APC-H7 | APC-H7 | | | Zombie NIR | Zombie NIR | | | | | | | |
| | | APC/Fire 810 | | | | | | | | | | | |

**Supplementary Table S1:** List of fluorochrome panels and instruments used. Abbreviations: “ID7000”: Sony ID7000™; “S8”: BD FACSDiscover™ S8; “NFB”: NovaFluor Blue; “NFY”: NovaFluor Yellow; “eF”: eFluor; “AF”: Alexa Fluor; “FVS”: Fixability Viability Stain; “Amth. Orange”: Amethyst Orange; “SuBr”: Super Bright.
